# Cargo recognition of Nesprin-2 by the dynein adapter Bicaudal D2 for a nuclear positioning pathway that is important for brain development

**DOI:** 10.1101/2025.05.18.654709

**Authors:** Estrella D Rodriguez Castro, Sivasankar Putta, M. Yusuf Ali, Jose M Garcia Martin, Xiaoxin Zhao, Samantha Sylvain, Kathleen M Trybus, Sozanne R Solmaz

## Abstract

Nesprin-2 and its paralog Nesprin-1 are subunits of LINC complexes that are essential for brain development. To position the nucleus for neuronal migration, Nesprin-2 interacts with the motors kinesin-1 and dynein, which are recruited by the adapter Bicaudal D2 (BicD2), but the molecular details of these interactions are elusive. Here, structural models of minimal Nesprin-2/BicD2 complexes with 1:2 and 2:2 stoichiometry were predicted using AlphaFold and experimentally validated by mutagenesis, binding assays and single-molecule biophysical studies. The core of the binding site is formed by spectrin repeats of Nesprin-2, which form an alpha-helical bundle with BicD2 that is structurally distinct from the Rab6/BicD2 and Nup358/BicD2 complexes. Such structural differences could fine tune motility of associated dynein and kinesin-1 motors for these transport pathways. Furthermore, the Nesprin-2 fragment interacts with full-length BicD2 and activates dynein/dynactin/BicD2 complexes for processive motility, suggesting that no additional components are required to reconstitute this transport pathway. Interestingly, either one or two Nesprin-2 molecules can bind to a BicD2 dimer and activate BicD2/dynein/dynactin complexes for processive motion, resulting in similar speed and run lengths. The BicD2/dynein-binding site is spatially close but does not overlap with the kinesin-1 recruitment site, thus both motors may interact with Nesprin-2 simultaneously. Several mutations of Nesprin-1 and 2 that cause Emery-Dreifuss muscular dystrophy are found in the motor-recruiting domain and may alter interactions with kinesin-1 and BicD2/dynein, consistent with the abnormally positioned nuclei found in patients with this disease.

## INTRODUCTION

Nesprin-2 has important roles in mechanotransduction and nuclear positioning during brain and muscle development, for which it recruits actin filaments and the motors dynein and kinesin-1 to the nuclear envelope ^1–4^. As a result, it mediates active crosstalk between the actin and microtubule networks ^5^. Embedded in the outer nuclear membrane, Nesprin-2 interacts with SUN (Sad1-UNC-84 homology domain) proteins in the inner nuclear membrane to form LINC complexes (linker of nucleoskeleton and cytoskeleton) ^6^. LINC complexes ^7–9^ span the nuclear envelope, interact with the lamina and chromatin ^10^ and act as force transducers between the nucleoskeleton and cytoskeleton.

The crucial role of Nesprin-2 (SYNE-2) and its paralog Nesprin-1 (SYNE-1) in brain development is evident from double-knockout mice, which die shortly after birth, due to severe defects in postmitotic neuronal migration and brain lamination during brain ^4^. Nesprin-2 alone appears to be essential for formation of the laminary structure in the hippocampus and cerebral cortex, whereas both Nesprin-1 and Nesprin-2 are important for the development of several other regions of the brain ^4^.

During neuronal migration, Nesprin-2 recruits the dynein adapter protein Bicaudal D2 (BicD2), which in turn recruits the minus-end directed microtubule motor cytoplasmic dynein ^2^. Nesprin-2 also has a LEWD sequence motif ^11^ that acts as a recruiting site for the plus-end directed microtubule motor kinesin-1 ^1,2,12,13^. During neuronal migration, dynein is the dominant motor which is actively modulated by kinesin-1. The association of motors with opposite polarity, such as dynein and kinesin-1, through adapter proteins is a common feature of cellular transport and likely fine-tunes motility, although the underlying molecular mechanisms remain unclear.

Nesprin-1 and Nesprin-2 are also important for the positioning of nuclei in developing myotubes, which are elongated, multinucleated muscle cells ^12,13^. In contrast to neuronal migration, kinesin-1 is the primary motor driving nuclear movement in myotubes, while dynein assists in this process ^12,13^. Depletion of either kinesin-1 or dynein motors or alternatively removal of the kinesin-1 recruiting LEWD motif of Nesprin-2 results in abnormal aggregation of the cell nuclei in the center of the myotube ^12,13^. Notably, several human disease mutations of Nesprin-2 and its paralog Nesprin-1 cause dilated cardiomyopathy, as well as Emery-Dreifuss muscular dystrophy (EDMD), which involves progressive skeletal muscle wasting and severe cardiomyopathy that often results in an early death ^14^. Abnormally clustered nuclei have been found in patients with autosomal dominant EDMD, suggesting that correct nuclear positioning may be required for proper muscle function. This highlights the important roles of Nesprin-2 in muscle development ^15^.

Nesprin-2 recruits actin filaments to its N-terminal calponin homology domains ^5,16,17^. The recruitment site for dynein/BicD2 and kinesin-1 was previously mapped to residues 6123 - 6421 of mouse Nesprin-2, which includes spectrin repeats (SR) SR52, SR53 and the adaptive domain (AD) ^1,2,13,18,19^. Subsequently, a mini-Nesprin-2 was constructed, which fused the N-terminal actin-recruiting domain to the microtubule motor recruitment domain located in SR52 - SR56, followed by the transmembrane alpha-helix and the C-terminal KASH (Klarsicht, Anc-1, Syne Homology) domain. While depletion of Nesprin-2 in rat embryo brains during development causes defects in neuronal migration, expression of the mini-Nesprin-2 restored near normal migration of neurons to the cortical plate. Deletion of the microtubule motor domain, however, resulted in a seven-fold increase of the distance between the nucleus and the centrosome and fewer neurons finished their migration and reached the cortical plate. These results suggest that the microtubule motor recruitment domain of Nesprin-2 has a key role in nuclear migration in neurons ^2^. In line with these results, the C-terminal truncation mutant BicD2 K774Ter reduces binding to Nesprin-2 and causes lissencephaly. This mutant results in reduced nuclear envelope localization of BicD2 and altered neuronal distribution in the neocortex of mice ^20^.

The involvement of Nesprin-1/2 in interkinetic nuclear migration, another brain developmental process, is debated. During this process, the nuclei of distinct brain progenitor cells move between the apical and basal brain surfaces, which is required for these cells to enter mitosis and undergo differentiation. This process appears to be largely unaffected by the removal of Nesprin-1 and Nesprin-2 from nuclear envelopes in brain progenitor cells with a dominant-negative approach ^21^, whereas a double knock-out approach results in interkinetic nuclear migration-related defects ^4,22^. In addition to the Nesprin-2-dependent nuclear positioning pathway, BicD2 facilitates two other transport pathways that are important for brain development. A second nuclear positioning pathway is driven by the recruitment of BicD2 to the nuclear pore protein Nup358 during the G2 phase of the cell cycle ^21,23^. This nuclear positioning pathway is essential for the differentiation of radial glial progenitor cells, which give rise to the majority of neurons and glia cells in the brain ^21^. Finally, BicD2 also interacts with Rab6^GTP^, a key effector of protein secretion, to recruit motors for the transport of secretory and Golgi-derived vesicles. Several of these secreted protein factors play critical roles in brain development ^24–29^. The importance of BicD2-dependent cellular transport pathways in brain and muscle development is underscored by the fact that human disease mutations of BicD2 cause devastating brain and muscle developmental defects, including spinal muscular atrophy, which is the most common genetic cause of death in infants ^18,30–32^. Several of the disease mutations cause changes in the affinity of BicD2 for distinct cargoes including Nup358, Nesprin-2 and Rab6^GTP^, which alters the associated cellular transport pathways ^18^. The cargo-binding domain (CTD) of BicD2 is located in the C-terminal portion of coiled coil 3 (CC3) ^26,28,33^. Rab6^GTP^, Nesprin-2 and Nup358 bind to distinct but overlapping binding sites on BicD2 and compete for binding, explaining the distinct effects of BicD2 mutations on the affinities towards these cargoes ^18,24,34–36^. This is also in line with the structural models of Rab6^GTP^/BicD2 and Nup358/BicD2, which we have previously established and validated ^24,34,36^. In addition, interactions of pathogens with BicD2 were characterized ^37,38^. However, a structural basis of how BicD2 recognizes its cargo adapter Nesprin-2 is elusive.

Here, we establish a structural model of the minimal Nesprin-2/BicD2 complex, which includes the minimal microtubule motor recruitment domain of Nesprin-2, using structure prediction by AlphaFold ^39^. This model was experimentally validated by mutagenesis and single-molecule functional assays. The core of the interaction is formed by SR52 and SR53 of Nesprin-2, which form a helical bundle with the BicD2-CTD. The BicD2/dynein binding site on Nesprin-2 is separated by a ∼65 residue disordered linker from the LEWD sequence motif that acts as a kinesin-1 binding site ^40^. Several mutations that cause Emery-Dreifuss muscular dystrophy are found in the motor-recruiting domain of Nesprin-1 and 2 ^41^ and may possibly alter interactions with kinesin-1 and BicD2/dynein, thereby modulating nuclear positioning pathways that are important for brain and muscle development.

## MATERIALS AND METHODS

### Structure predictions

The structure predictions were carried out using ColabFold v1.5.5 ^42^ with the Google Colab AlphaFold2_mmseq2 Notebook (https://colab.research.google.com/github/sokrypton/ColabFold/blob/main/AlphaFold2.ipynb, accessed on 19 March 2025), which implements AlphaFold ^39,42,43^. Default settings were used; a template was not applied. A 2:2 hetero-dimer composed of two molecules of BicD2-CTD (human BicD2 aa 715 – 804, which is identical in sequence to mouse BicD2 aa 711-800) and two molecules of mouse Nesprin-2 (Uniprot ID Q6ZWQ0) (aa 6123 – 6421)^1,18,40,42^ was predicted. We also predicted a 1:2 complex with a single Nesprin-2 molecule. The prediction of the larger 2:2 complex includes the sequence of human BicD2 (aa 667 - 804) and mouse Nesprin-2 (aa 5692 - 6342). The predicted structure of the minimal Nesprin-1/BicD2-CTD complex includes human Nesprin-1 (Uniprot ID Q8NF91) aa 7998 – 8315 and BicD2 aa 715 - 804. UCSF ChimeraX was used to create structure figures ^44^. The PISA server was used to identify contact residues between Nesprin-2 and BicD2 ^45^.

### GST-pulldown assay

All expression vectors were obtained from the company Genscript that performed gene synthesis of codon-optimized inserts, cloning and mutagenesis. Expression constructs of N-terminally GST-tagged mouse Nesprin-2 (aa 6123 – 6421, Uniprot ID Q6ZWQ0) in the pGEX6p1 expression vector and N-terminally his_6_-tagged human BicD2-CTD (aa 715 – 804; this fragment is identical in sequence to mouse BicD2-CTD aa 711-800) in the pET28a expression vector were previously described ^18,46,47^. Human Nesprin-1 (Uniprot ID Q8NF91) aa 7997 – 8328 was cloned into the same vector. Mutagenesis was performed by Genscript. These protein fragments were expressed in *E. coli* and purified as previously described ^18^. The BicD2 fragment was expressed in the LOBSTR BL21(DE3)-RIL strain at 37 °C for three hours, and the Nesprin-1/2 fragments were expressed in the BL21(DE3)- RIL strain at 16°C for 12-20 h.

GST pulldowns with GST-tagged Nesprin-2 and BicD2-CTD fragments (wild-type (WT), mutants or deletions) were performed as described ^18^. His_6_-tagged BicD2-CTD (WT or mutants) was purified from 1L of cell culture by Ni-NTA affinity chromatography, using a salt concentration of 250 mM NaCl. The purified protein was analyzed by SDS-PAGE and diluted to reduce the salt concentration to 125 mM. GST-tagged Nesprin-1 or Nesprin-2 fragments were purified from 0.5 L of cell culture by glutathione affinity chromatography but not eluted (a cOmplete EDTA-free protease inhibitor cocktail tablet (Roche) was added to the Nesprin-1 lysate during purification). The column was incubated for 30 min on a nutator with the purified BicD2-CTD, and the column was washed twice with binding buffer and eluted with 10 mM glutathione, 50 mM Tris pH 8.0, 150 mM NaCl, 1 mM DTT. Elution fractions were analyzed by SDS-PAGE using 16% acrylamide gels and stained with Coomassie Blue. The background subtracted intensities of the gel bands corresponding to the Nesprin-2 and BicD2 fragments were quantified using ImageJ ^48^ as described by ^18^. The ratio of bound BicD2-CTD/Nesprin-2 was calculated and normalized to the WT (WT = 1) positive control, which was analyzed on the same SDS-PAGE as the mutant.

For Figure 2, the intensities of the distinct Nesprin-2 and BicD2 bands were divided by the molecular weight prior to calculating the ratio of bound BicD2/Nesprin-2 and the average ratio for the WT (WT=1) was used for normalization.

### Circular Dichroism (CD) spectroscopy

CD spectroscopy was performed as described ^24^. For these experiments, GST-tagged Nesprin-2 aa 6123 – 6421 fragments (WT and mutants) were expressed and purified as described ^18^ with the GST-tag intact. For Figure S12, the Nesprin-2 fragment was purified by glutathione affinity chromatography and eluted by proteolytic cleavage with 500 Units of PreScission protease (GE Healthcare) per L of cell culture for 16 h. The purified proteins were transferred into a buffer 10 mM Tris, pH 8.0, 150 mM NaCl, 0.2 mM TCEP (Tris(2-carboxyethyl)phosphine) by three cycles of dilution and concentration in a centrifugal concentration filter as described ^24^. The protein concentration was 0.5 mg/ml unless otherwise noted. The protein concentration was determined by absorbance spectroscopy at 228.5 nm and 234 nm using the extinction coefficient of the peptide bond and flash-frozen in liquid nitrogen as described ^36^.

CD measurements were recorded at a temperature of 10°C with the use of Jasco J-1100 CD spectrometer including CD, HT, and absorbance signals at a wavelength range of 250 to 190 nm. A quartz cuvette with a pathlength of 1 mm was used. The following parameters were used: data pitch: 0.1 nm; D.I.T.: 2 s; bandwidth: 1.00 nm; scanning speed: 50 nm/min; 8 accumulations per sample. The wavelength scan of the buffer was subtracted from the recorded CD wavelength scans, and the raw ellipticity Θ (mdeg) was converted to mean residue molar ellipticity [Θ] (path length: 1 mm; number of peptide bonds: 542 (Figure 4); or 426 (Figure S12); WT molar mass: 63,531.5 Da (Figure 4); or 49,100 Da (Figure S12)). For Figure 4, the protein concentration used for the calculation was determined from the absorbance measurement from the CD instrument at 228.5 nm and 234 nm using the extinction coefficient of the peptide bond ^36^, after subtracting the buffer baseline. For Figure S12, the protein concentration for the calculation was determined from the absorbance measurement at 214 nm using the extinction coefficient (124168 M-1 cm-1) at 214 nm calculated by BeStSel, after subtracting the buffer baseline. BeStSel was used to estimate the secondary structure from the CD wavelength scans (200 – 250 nm)^49^. For each protein, three CD wavelength scans were recorded from separate purifications, and a representative result is shown.

### Protein expression and purification for single molecule assays

Cytoplasmic dynein and microtubules were purified from tissue (bovine brain) according to established protocols ^50^. Specifically, dynein and dynactin were isolated from 300 g of bovine brain following the procedure of ^51^, while tubulin was purified from 200 g of bovine brain as described previously ^50^. The Uniprot ID for human Bicaudal D2 is NP_001003800.1. The N-terminal domain of human BicD2 (BicD2^CC1^), containing an N-terminal biotin tag for labeling purposes, was expressed in *E. coli* and purified as described ^50^. Full-length, wild-type human BicD2 was expressed in Sf9 cells and purified using the same protocol as for *Drosophila* BicD ^36^. Full-length mouse kinesin-1, including both heavy and light chains (Uniprot ID NP_032474.2), was expressed with a C-terminal biotin tag and purified as described for the *Drosophila* homolog^52^. Nesprin-2 fragments (for protein sequences see Figure S13) were expressed and purified as N-terminal GST fusion proteins as described ^18^, using glutathione affinity chromatography and size exclusion chromatography with the Superdex 200 Increase 10/300 GL column (GE Healthcare), which was equilibrated with the following buffer: 20 mM HEPES pH 7.4, 150 mM NaCl, 0.5 mM TCEP. For Nesprin-2 aa 6123-6421, purification was performed with a SNAP tag (Figure S13). The SNAP tag was converted to SNAP-biotin using SNAP-biotin (New England Biolabs, Catalog #S9110S), enabling binding to Alexa Fluor 488–streptavidin ^53^. Nesprin-2 aa 6123-6352 WT and the V6246A/L6253A/F6261A triple mutant were purified with biotin tags at their N-terminal domain (Figure S13). Protein concentrations for all protein preparations were measured using the Bradford assay (Bio-Rad).

### Single molecule binding assays with Nesprin-2, full-length BicD2 and kinesin-1

To investigate the interaction between BicD2 and Nesprin-2, or kinesin-1 and Nesprin-2, BicD2, Nesprin-2 fragments, and kinesin-1 were clarified by ultracentrifugation at 400,000 × g for 20 min at 4 °C to remove aggregates and then diluted to 2 µM in BRB80 buffer (80 mM PIPES, 1 mM MgCl₂, 1 mM EGTA) supplemented with 20 mM DTT (dithiothreitol). For the Nesprin-2–kinesin-1 interaction, biotin-tagged Nesprin-2 (2 µM) was mixed with streptavidin-coated Alexa Fluor 488 (2 µM), and biotin-tagged kinesin-1 (2 µM) was mixed with streptavidin-coated Alexa Fluor 647 (2 µM) at a 1:1 molar ratio. Samples were incubated for 15 min to allow labeling. The labeled kinesin-1 and Nesprin-2 fragments were then combined and incubated on ice for 30 min to form kinesin-1/Nesprin-2 complexes. Similarly, for the BicD2–Nesprin-2 interaction, BicD2 and Nesprin-2 were labeled with Alexa Fluor 647–streptavidin and streptavidin–Alexa Fluor 488, respectively, and processed as described above.

The complexes were diluted 100-fold (final concentration: 5 nM) in BRB80 buffer. Samples were introduced onto the glass surface and incubated for 3 min, followed by washing with BRB80 buffer. To prevent photobleaching, an oxygen scavenger system consisting of 5.8 mg/mL glucose (EM Science, DX0145), 0.045 mg/mL catalase (Sigma-Aldrich, C40), and 0.067 mg/mL glucose oxidase (Sigma-Aldrich, G6125) was included in the BRB80 buffer.

Images were acquired using a total internal reflection fluorescence (TIRF) microscope equipped with two cameras for simultaneous dual-color imaging. Colocalization of the two differently labeled proteins was identified by the appearance of yellow fluorescence (signal overlap).

#### Single molecule processivity assays

Dynein, dynactin, BicD2, and Nesprin-2 constructs were diluted in BRB80 buffer supplemented with 20 mM DTT and clarified to remove aggregates as described above. To assemble the dynein–dynactin–BicD2–Nesprin-2 (DDBN) complex, components were mixed at a 1:1:1:2 molar ratio (250 nM dynein, 250 nM dynactin, 250 nM BicD2, and 500 nM Nesprin-2) and incubated at room temperature for 30 min. Similarly, dynein–dynactin–BicD2 (DDB) and dynein–dynactin–BicD2^CC1^ (DDB^CC1^) complexes were reconstituted at a 1:1:1 molar ratio (250 nM dynein, 250 nM dynactin, and 250 nM BicD2 or BicD2^CC1^).

Complexes containing biotinylated BicD2 were labeled with streptavidin-conjugated Alexa Fluor 488 Qdots at a 1:1 molar ratio (BicD2:Qdot). This ratio was chosen to minimize binding of multiple BicD2 molecules to a single streptavidin-Qdot. In the motility assay, to directly compare the three complexes (DDB, DDBN, and DDB^CC1^), only BicD2 was fluorescently labeled, while dynein, dynactin, and Nesprin-2 remained unlabeled. The labeled complexes were further diluted into motility buffer (BRB80 supplemented with 20 mM DTT, 5 mg/ml BSA (bovine serum albumin), 0.5 mg/ml κ-casein, 0.5% Pluronic F-68, 10 μM paclitaxel, and an oxygen scavenging system) to a final concentration of 5 nM dynein for motility assays. The oxygen scavenging system was added immediately before imaging. Samples were incubated for 15 min at room temperature prior to observation by TIRF microscopy.

To determine the number of Nesprin-2 molecules bound to DDBN, equal molar amounts of Nesprin-2 fragment were mixed with either 525-nm or 655-nm streptavidin–Qdots in two separate microcentrifuge tubes and incubated for 15 min. To block the unoccupied binding sites on the Qdots, 5 µM biotin were added to both tubes and incubated for an additional 10 min. Equal amounts of 525-nm– and 655-nm–labeled Nesprin-2 fragments were subsequently added to dynein, dynactin, BicD2 (DDB). The final molar ratio of the dynein–dynactin–BicD2–Nesprin-2 (DDBN) complex was 1:1:1:2 (250 nM dynein, 250 nM dynactin, 250 nM BicD2, and 500 nM Nesprin-2). The mixture was incubated for 30 min to form the DDBN complex. In this experiment, dynein, dynactin, and BicD2 were unlabeled; only Nesprin-2 was labeled with either green or red Qdots to determine whether one or two Nesprin-2 molecules were bound to BicD2 for the processive motion of DDBN. The reconstituted complexes were diluted to 5 nM dynein in BRB80 buffer and introduced into a flow chamber containing surface-immobilized microtubules as described above.

Rhodamine-labeled microtubules and PEGylated glass slides for motility assays were prepared according to established protocols ^50^. To facilitate microtubule attachment, slides were first coated with 0.5 mg/ml rigor kinesin-1, washed 2–3 times with motility buffer, and then incubated with rhodamine-labeled microtubules. Excess microtubules were removed by 2–3 additional washes with motility buffer. Finally, reconstituted complexes (DDB, DDB^CC1^, or DDBN) were introduced into the flow chambers for imaging.

To observe the motility of Qdot-labeled DDB, DDB^CC1^, and DDBN complexes on microtubule tracks, we performed TIRF microscopy as described previously ^52^. Imaging was carried out on a Nikon ECLIPSE Ti microscope with an objective-type TIRF system and controlled by Nikon NIS-Elements software. Two Andor iXon Ultra EMCCD cameras (Andor Technology USA, South Windsor, CT) were used to capture moving complexes. To detect the dual-color complex on the glass surfaces, 30 – 60 frames were captured at 200 ms intervals. Single-molecule images were collected using Alexa Fluor 488 or 647 and a 561-nm laser was used for rhodamine-labeled microtubules. Alexa Fluor 488 was excited using the 488 nm laser line, with excitation and emission peaks at 496 nm and 519 nm, respectively. Alexa Fluor 647 was excited with the 640 nm laser line, exhibiting excitation and emission peaks at 650 nm and 671 nm. Both quantum dots (525 nm and 655 nm) were excited using the 488 nm laser line, emitting at 525 nm and 655 nm, respectively. A GFP/RFP dichroic filter was used to efficiently separate excitation and emission by reflecting the excitation wavelengths and transmitting the fluorescence signals from both GFP and RFP. Movies consisted of 300 - 600 frames acquired at 100 - 200 ms intervals (10 - 5 frames/s) using two Andor EMCCD cameras (Andor Technology USA, South Windsor, CT). Images were acquired at a spatial resolution of 0.1066 μm per pixel. Individual Qdots were tracked with the ImageJ 1.54g MTrackJ plugin ^54^ to measure the run length of single complexes. Speed was calculated by dividing the total distance traveled by a single molecule by the total time. A processive event was defined as movement greater than 0.3 µm.

Run length distributions were plotted as 1 – cumulative probability distribution (1 – CDF) in GraphPad Prism 10, and characteristic run lengths were determined by fitting the data to a one-phase exponential decay equation: p(x)=Ae^−x/λ^ where p(x)is the relative frequency, x represents the travel distance along the microtubule, and A denotes the amplitude ^36,55^. For speed, the mean values were determined using GraphPad Prism.

The binding rate was quantified by counting Qdot-labeled complexes bound to microtubules per unit time and per μm of microtubule.

Comparisons between two datasets, such as run length, were assessed using the Kolmogorov–Smirnov test, whereas statistical significance among three or more datasets, including binding rate, or binding assays, was determined by the one-way ANOVA followed by Tukey’s post-hoc test.

## RESULTS

### A structural model of the minimal Nesprin-2/BicD2 complex with a PAE score in the high confidence range was obtained using AlphaFold

BicD2 recruits dynein, dynactin and kinesin to the cargo adapter Nesprin-2, for a nuclear positioning pathway that is important for muscle development and neuronal migration, a fundamental process in brain development ^2^. We previously reconstituted a recombinant minimal Nesprin-2/BicD2 complex ^18^, with the C-terminal cargo binding domain (CTD) of human BicD2 (aa 715 - 804) and spectrin repeats (SR) SR52 and SR53 of Nesprin-2, as well as the AD (adaptive) domain (aa 6123 to 6421). This Nesprin-2 domain has been previously mapped as the minimal recruiting domain for dynein and kinesin-1 ^1,2,5,13,18,56^.

Knowledge of the stoichiometry of the complex is a key parameter for the structure prediction. Therefore, we characterized the oligomeric state of a purified minimal Nesprin-2/BicD2 complex by size exclusion chromatography coupled to multi-angle light scattering (SEC-MALS), which allows determination of the molar mass across a size exclusion elution peak with high accuracy (5% error) (Figure S1). The weight-averaged molar mass for the first elution peak is MW= MW=128.9 ± 4.9 kDa, which closely matches the calculated mass of a 2:2 complex of Nesprin-2 and BicD2 (130.4 kDa; note that the mass of a BicD2 protomer is 10.9 kDa, and the mass of a Nesprin-2 protomer is 54.3 kDa). It should be noted that the GST-tag may affect the oligomeric state, and we have included functional data (see below), which suggest that Nesprin-2/BicD2 complexes with both a 1:2 and 2:2 stoichiometry are capable of activating dynein/dynactin for processive motion. It should be also noted that two other BicD2/cargo complexes (Nup358/BicD2 and Rab6^GTP^/BicD2, without affinity tags such as GST) were previously characterized by SEC-MALS, which established a 2:2 stoichiometry ^35,36,46^. Furthermore, the X-ray structure of the human BicD2-CTD was previously determined, which confirmed that it forms a homodimer ^35,47^. To establish a structural basis for the interaction of the dynein adapter BicD2 with Nesprin-2, we predicted the structure of the minimal 2:2 complex with ColabFold ^42^, which implements the AI-based structure prediction software AlphaFold ^39,43^.

The structural model of this minimal Nesprin-2/BicD2 complex with the lowest predicted aligned error (PAE) is shown in Figure 1, and the remaining predicted models, which are overall quite similar, are shown in Figure S2. In the structural model, SR52 and SR53 of Nesprin-2 form a helical bundle with the BicD2-CTD. However, the intrinsically disordered AD domain, which includes the kinesin-1-recruiting LEWD motif does not engage in the interaction with BicD2.

**Figure 1.**
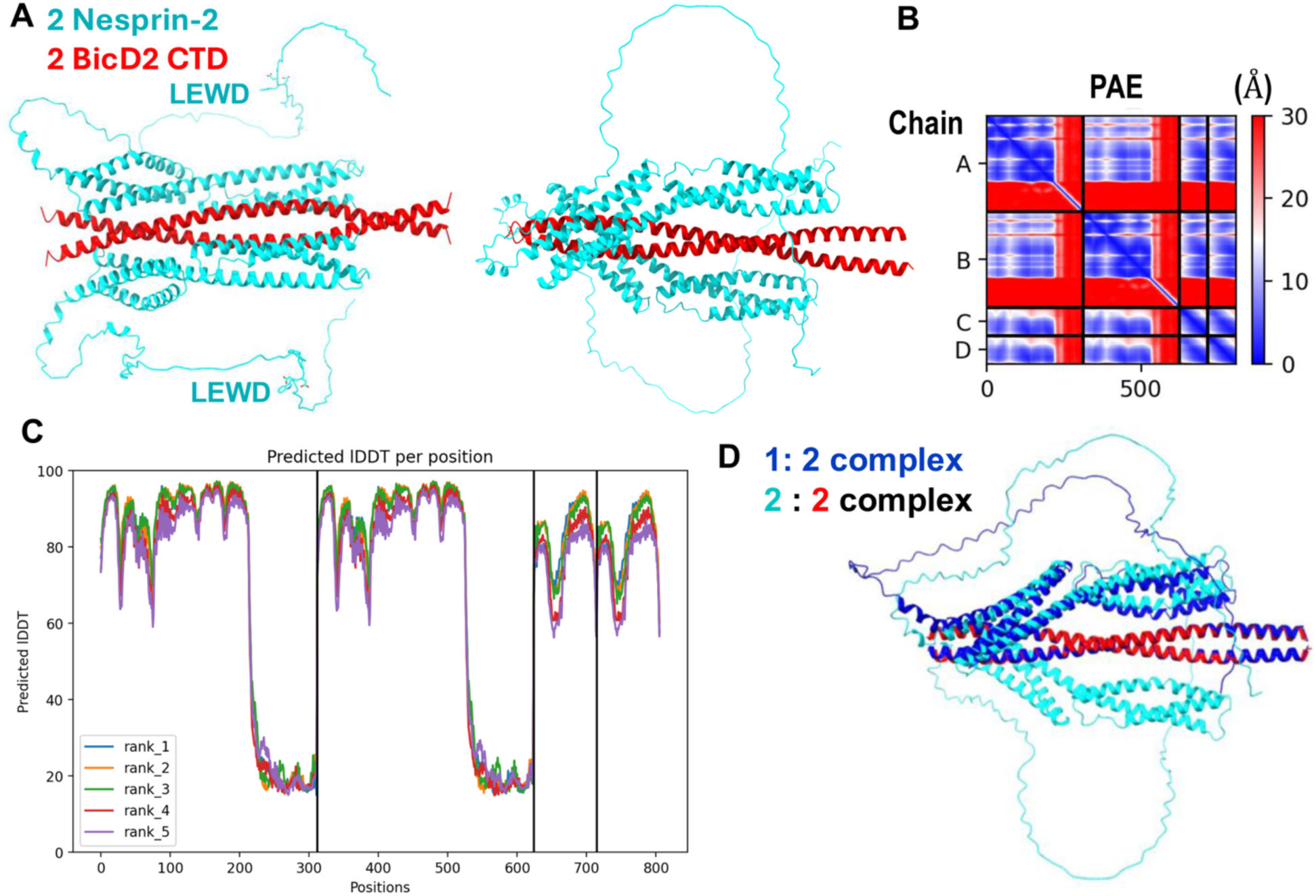
A structural model of the minimal Nesprin-2/BicD2 complex with a reliable PAE score was obtained from AlphaFold. (A) Structural model of the minimal Nesprin-2/BicD2 complex (Nesprin-2-SR52-SR53AD and BicD2-CTD) from AlphaFold, shown in cartoon representation, rotated by 90°. The LEWD sequence motifs, which act as kinesin-1 binding site are labeled and shown in stick representation. (B) Predicted aligned error (PAE) plot for the highest-ranking model. The x and y axes show the residue numbers, starting with Nesprin-2 (aa 6123 – 6421, Chain A, B), followed by BicD2 (aa 715-804; chain C, D) ^1,18,57^. (C) pLDDT error plot. (D) The AlphaFold prediction of the Nesprin-2/BicD2 complex with 1:2 stoichiometry (blue) is least-squares superimposed with the 2:2 complex (cyan and red). Additional models and error plots are shown in Figures S2, S3.

A key metric for evaluation of the prediction is the PAE, as it provides an error estimate for the distances of each residue pair. The PAE of the highest-ranking model is mostly below 5 - 10 Å, reflecting a high degree of confidence in the predicted structure (Figure 1B) ^43^, especially considering that an experimental structure of the Nesprin-2 fragment or a homolog is not available. The only region with a higher PAE error is the AD domain, which is to be expected, as it is intrinsically disordered, resulting in a larger variability of inter-residue distances. This region does not engage in the interaction with BicD2-CTD. Overall, the PAE of the predicted Nesprin-2/BicD2 interface is comparable to the PAE of the previously characterized predicted Nup358/BicD2 interface ^34^. The PAE of the predicted Rab6^GTP^/BicD2 interface was lower, likely because an experimental structure of Rab6^GTP^ is available ^24^.

The predicted local distance difference test (pLDDT) ^39,58^ is a per-residue metric of local confidence. It estimates how well the predicted structure would agree with an experimental structure based on a local distance difference test. The pLDDT scores for all predicted models span between 80 - 98, suggesting that they are reliable (based on the established 70 confidence threshold ^58^, and the pLDDT patterns are similar for the five top-ranking models (Figure 1C). It should be noted that the intrinsically disordered AD domain of Nesprin-2 that does not interact with BicD2, and the residues in the loop regions that connect the alpha-helices of the spectrin repeats have low pLDDT scores, as expected for disordered domains (Figure 1C, S2).

Furthermore, we also used AlphaFold to predict the structure of a Nesprin-2/BicD2 complex with a 1:2 stoichiometry (Figure 1D, S3), to assess if complexes with alternative stoichiometry may be formed. The resulting structure closely resembles the structure of the 2:2 complex, with only one of the two Nesprin-2 binding sites occupied (Figure 1D, S3). The root mean square deviation (RMSD) of the superimposition of the 1:2 and 2:2 complex is 1.8 Å, which suggests close similarity of the structures, and the PAE and pLDDT plots are overall similar for both predictions (Figure S3).

Nesprin-1 is a paralog with overall similar functions as Nesprin-2. Interestingly, several human disease mutations of Nesprin-1 cause Emery-Dreifuss muscular dystrophy ^41^, but it has not been established whether Nesprin-1 interacts with BicD2 as well. The human Nesprin-1 domain that includes SR70, SR71 and a disordered domain with a LEWD motif (aa 7998 - 8315) has 93% sequence conservation with the Nesprin-2-SR52-SR53AD domain (Figure S4). To confirm the interaction between this Nesprin-1-SR70SR71AD domain and BicD2-CTD, we performed a GST (Glutathione-S-Transferase) pulldown-assay with recombinantly expressed proteins (Figure S4). The Nesprin-1 fragment and BicD2 co-elute, suggesting a direct interaction between Nesprin-1 and BicD2. A structural model of this minimal Nesprin-1/BicD2 complex was obtained from Alphafold-2, which has PAE and pLDDT scores in the reliable range and closely resembles the structural model obtained for the minimal Nesprin-2/BicD2 complex (Figure S5). These results suggest that Nesprin-1 may also interact with the BicD2-CTD domain.

Furthermore, we predicted the structure of a larger 2:2 complex including SR48 - SR53 of Nesprin-2 (aa 5692 - 6342) and the entire coiled coil 3 domain of human BicD2 (aa 667-804). This larger complex was chosen based on recent results suggesting that inclusion of SR 48 increased the interaction of Nesprin-2 with BicD2 based on qualitative binding assays ^40^.

Figure S6 shows the highest-ranking model from the structural prediction of this larger complex. A least square superimposition of the structural models of the larger complex with Nesprin-2 SR48 – SR53 and BicD2-CC3 and the minimal Nesprin-2-min/BicD2-CTD complex suggests that they are very similar in the shared domains, supporting that the structural model of the minimal complex is reliable, since the addition of the additional regions does not significantly change the structure prediction of the minimal complex. However, the PAE plot of the larger complex indicates a high error for the additional domains, suggesting that the structure prediction of the added domains is not reliable. Due to the high PAE errors and the differences of the predictions (Figure S7), we concluded that only the structure prediction of the minimal Nesprin-2/BicD2 complex was reliable, because its PAE is mostly below 5 - 10 Å. Thus, we focused in the subsequent structural analysis on this complex. It should be noted that additional, weaker interfaces between Nesprin-2 and BicD2 may be formed in the context of the full-length proteins.

### Validation of the structural model of the Nesprin-2/BicD2-CTD complex

To validate the structural model of the minimal Nesprin-2/BicD2-CTD complex, we designed a series of deletion constructs of Nesprin-2 and assessed binding to BicD2-CTD by GST-pulldown assays (Figure 2). The minimal Nesprin-2 construct (aa 6123 - 6340) that still interacts robustly with BicD2-CTD in the pulldown assays lacked the entire intrinsically disordered domain (Figure 2), confirming that it is dispensable for BicD2-CTD binding, consistent with our structural model. Of note, this domain includes the LEWD sequence motif that acts as a binding site for the kinesin-1 light chain 2 (Figure 2), suggesting that BicD2/dynein and kinesin-1 could interact simultaneously with Nesprin-2. An additional N-terminal deletion construct was designed to remove SR52 and SR53. As expected, this construct did not pull down visible amounts of BicD2-CTD (Figure 2).

**Figure 2.**
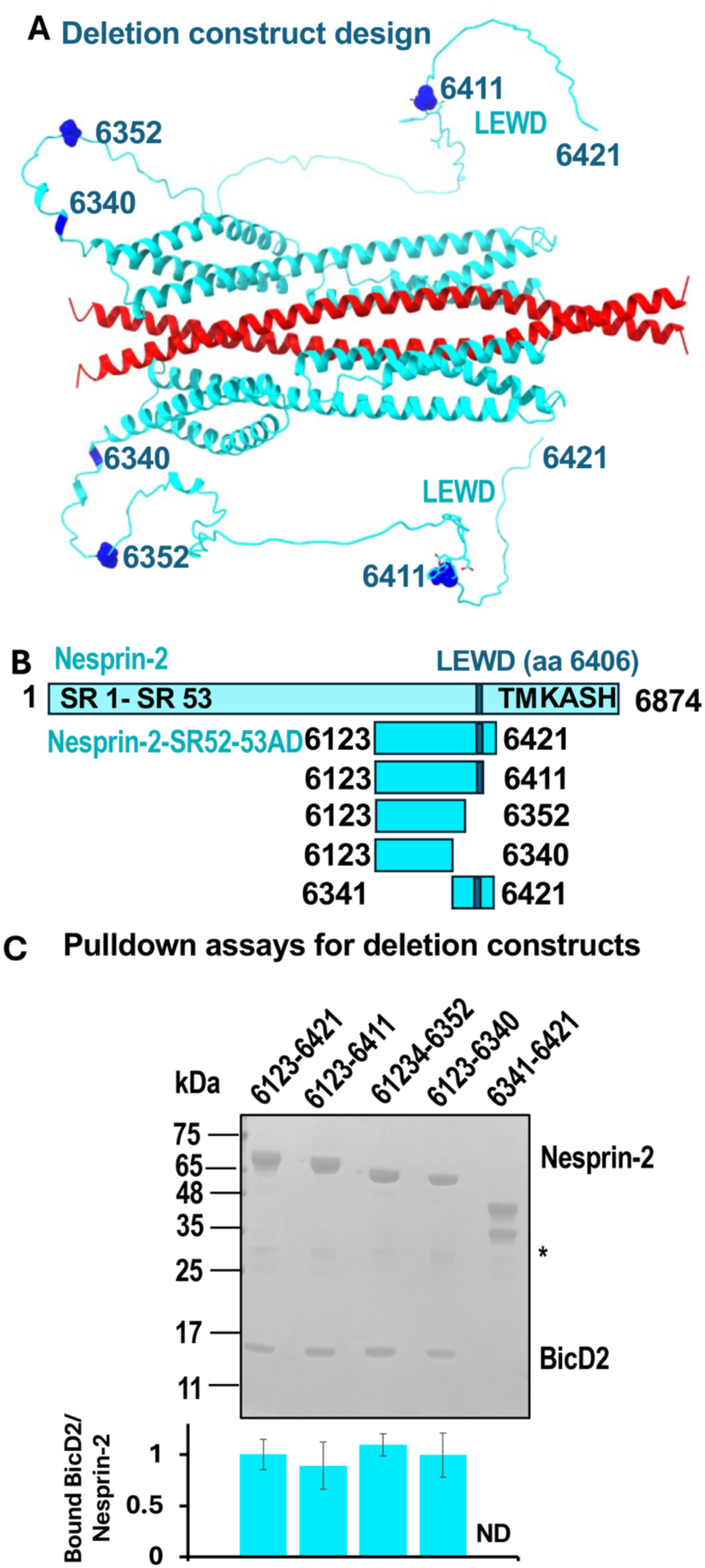
The intrinsically disordered domain with the LEWD sequence motif that acts as kinesin-1 binding site is dispensable for the interaction with BicD2. (A) Structural model of the minimal Nesprin-2/BicD2 complex. The residue numbers for deletion sites are highlighted in dark blue and labelled. The LEWD motif is shown in stick representation. (B) Schematic representation of the deletion constructs. (C) Pulldown assays were conducted with the GST-tagged Nesprin-2 deletion constructs from B and BicD2-CTD. The SDS-PAGE of the elution fractions is shown, and the quantification for each gel lane is shown below as a bar graph. The ratio of bound His_6_-tagged BicD2-CTD per GST-tagged Nesprin-2 fragment was quantified from the intensity of the gel bands and normalized respective to the average ratio of Nesprin-2 (aa 6123- 6421)/BicD2-CTD = 1. Data were averaged from at least 3 replicates; the standard deviation is shown. Note that all C-terminal deletions still interact with BicD2-CTD, suggesting that the intrinsically disordered regions with the LEWD motif are dispensable for the interaction. The asterisk indicates GST. ND: not determined. A negative control is shown in Figure S9.

To further validate our structural model of the minimal Nesprin-2/BicD2-CTD complex, we mutated contact residues of Nesprin-2 and BicD2 (Table S1) from the structural model to alanine. The interaction between Nesprin-2 and BicD2 (WT or mutants) was assessed by GST-pulldown assays (Figure 3). Five mutants of BicD2-CTD that reduced binding to Nesprin-2 were identified (red in Figure 3). These mutants were located in the extreme C-terminus of the BicD2-CTD, which was previously mapped as the minimal Nesprin-2 binding site ^18^. Notably, five Nesprin-2 mutants (V6246, E6250, H6260, Y6319, and R6329) were identified that displayed reduced binding to BicD2-CTD (red in Figure 3). Several additional Nesprin-2 mutants had reduced expression and were excluded from the analysis as they may be misfolded (Figure S9A). Of note, a triple mutant (V6246A/L6253A/F6261A) was designed to disrupt three key hydrophobic contacts between Nesprin-2 and BicD2 that span most of the SR53 (Figure S8). This mutant virtually abolishes the interaction between Nesprin-2 and BicD2-CTD in pulldown assays and was chosen for further characterization.

**Figure 3.**
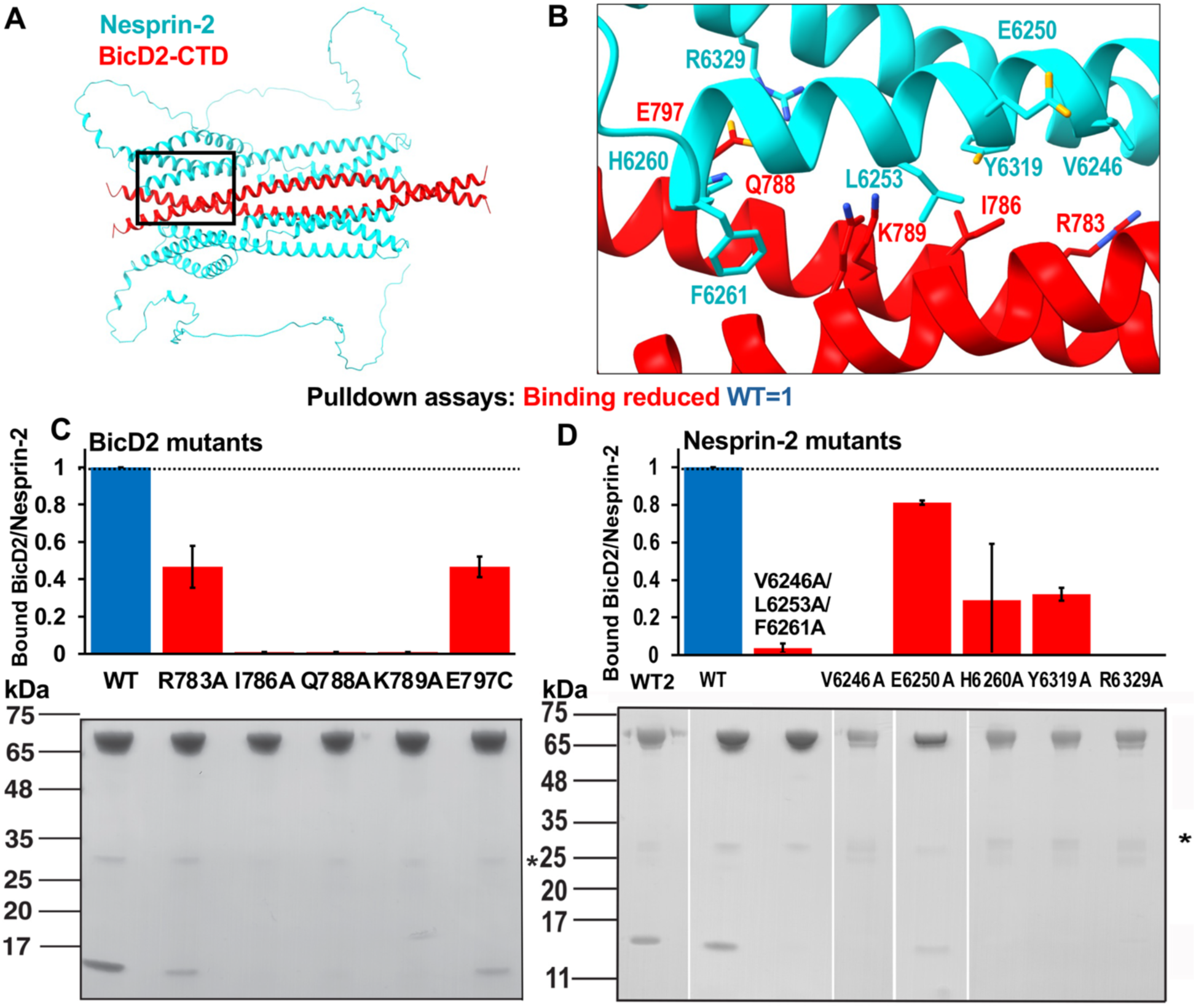
The structural model of the minimal Nesprin-2/BicD2 complex was validated by mutagenesis and binding assays. (A) Structural model of the minimal Nesprin-2/BicD2 complex. The boxed region is enlarged in B. (B) The Nesprin-2 (cyan) and BicD2 (red) contact residues from the AlphaFold model that were validated by mutagenesis in C and D are shown in stick representation and labelled. (C) BicD2 residues that formed contacts with Nesprin-2 in the structural model were mutated to alanine and the interaction was probed by GST-pulldown-assays. The elution fractions were analyzed by SDS-PAGE and the ratio of bound His_6_-tagged BicD2-CTD per GST-tagged Nesprin-2 (aa 6123-6421) was quantified from the intensity of the gel bands and normalized respective to the WT (WT=1). Bar graphs of the ratio of bound BicD2/Nesprin-2 from three experiments (n=3) and the standard deviation are shown (blue: WT; red: mutants with reduced binding). The SDS-PAGE of the elution fractions of one respective experiment is shown below the bar graph. A loading control and a negative control for the pulldown (i.e. without a GST-tagged protein) are shown in Figure S9A. (D) Nesprin-2 residues that formed contacts with BicD2-CTD were mutated and analyzed as described in C (n=3). WT is the wild-type control for the triple mutant and E6250A; WT2 is the wild-type control for all other mutants. The asterisk indicates GST.

Figure 3B shows a closeup of the interface between Nesprin-2 and BicD2-CTD, in which the residues that were confirmed to be important for the interaction by pulldown assays are shown in stick representation and labelled. Most of the confirmed Nesprin-2 contact residues are hydrophobic, whereas some of the BicD2 contact residues are also charged.

To exclude that these mutations result in structural changes or misfolding of Nesprin-2, which could also result in reduced binding, we assessed the secondary structure of Nesprin-2 and the mutants by circular dichroism (CD) spectroscopy (Figure 4). Two local minima (at 208 nm and 222 nm) are observed for WT Nesprin-2, which are characteristic for alpha-helical structures. The wavelength scans of six Nesprin-2 mutants with reduced binding to BicD2 (see Figure 3D) are very similar to the wavelength scan of WT Nesprin-2 (the experimental error of the molar ellipticity is 3.5 - 5% ^36,59^), suggesting that the mutations do not result in misfolding or large structural changes compared with the WT. These results validate the mutated residues with reduced binding as bona fide contact residues.

**Figure 4.**
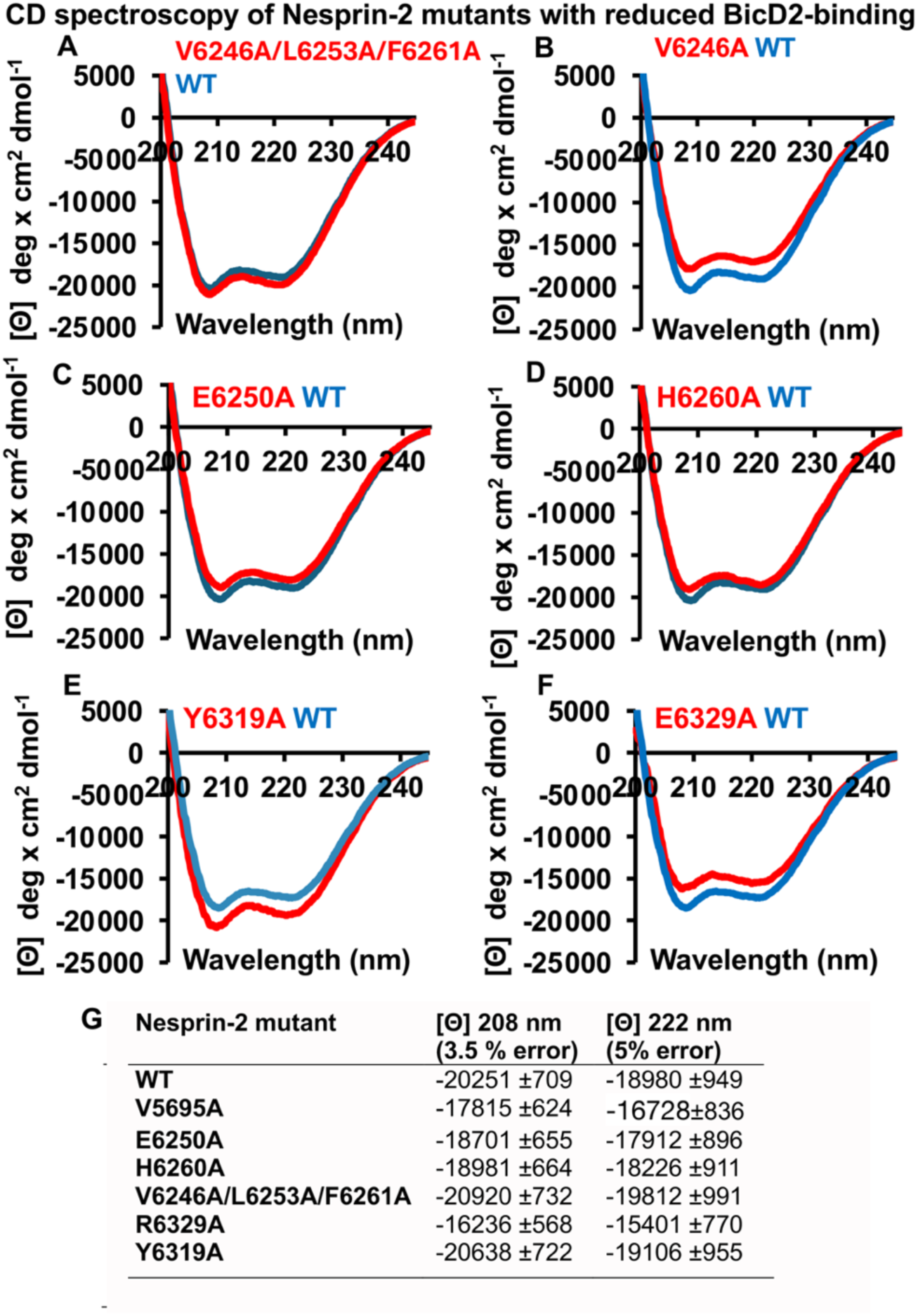
Circular dichroism (CD) wavelength scans of Nesprin-2 mutants with reduced BicD2 binding are comparable to wild-type (WT) spectra, suggesting the mutations do not induce major structural changes. (A-F) The CD wavelength scans of six purified GST-tagged Nesprin-2 (aa 6123-6421) mutants that reduce binding to BicD2 from Figure 3 D are shown in red and overlaid with the CD wavelength scan of the WT (blue). The molar ellipticity [8] is plotted versus the wavelength. The experiment was repeated three times, and a representative experiment is shown. (G) Molar ellipticity values for all Nesprin-2 fragments at two wavelengths that are characteristic for alpha-helical structures: 208 and 222 nm. The experimental error at these wavelengths was previously determined ^59^. Table S2 shows the secondary structure estimation from the CD spectra by the program BeStSel ^49^.

Notably, mutagenesis confirmed several residues to be essential for the Nesprin-2/BicD2 interaction, thereby validating our structural model of the minimal Nesprin-2/BicD2-CTD complex. The structural model was also validated by deletion constructs which established that the intrinsically disordered AD domain is dispensable for the interaction, in line with our structural model.

### Comparison of the structural models of three BicD2/cargo complexes that are important for brain development

In addition to the structural model of the minimal Nesprin-2/BicD2 complex presented here, structural models for minimal Nup358/BicD2 and Rab6/BicD2 complexes were previously obtained from AlphaFold and validated through experiments ^24,34,36^. These experiments suggest that the three cargoes Nesprin-2, Nup358 and Rab6^GTP^ bind to distinct but overlapping binding sites on BicD2, consistent with our previous results that these cargoes compete for binding ^18,35^, and consistent with the distinct effects of human BicD2 disease mutations on the affinity of different cargoes (Figure 5, Figure S10, Figure S11) ^18^.

**Figure 5.**
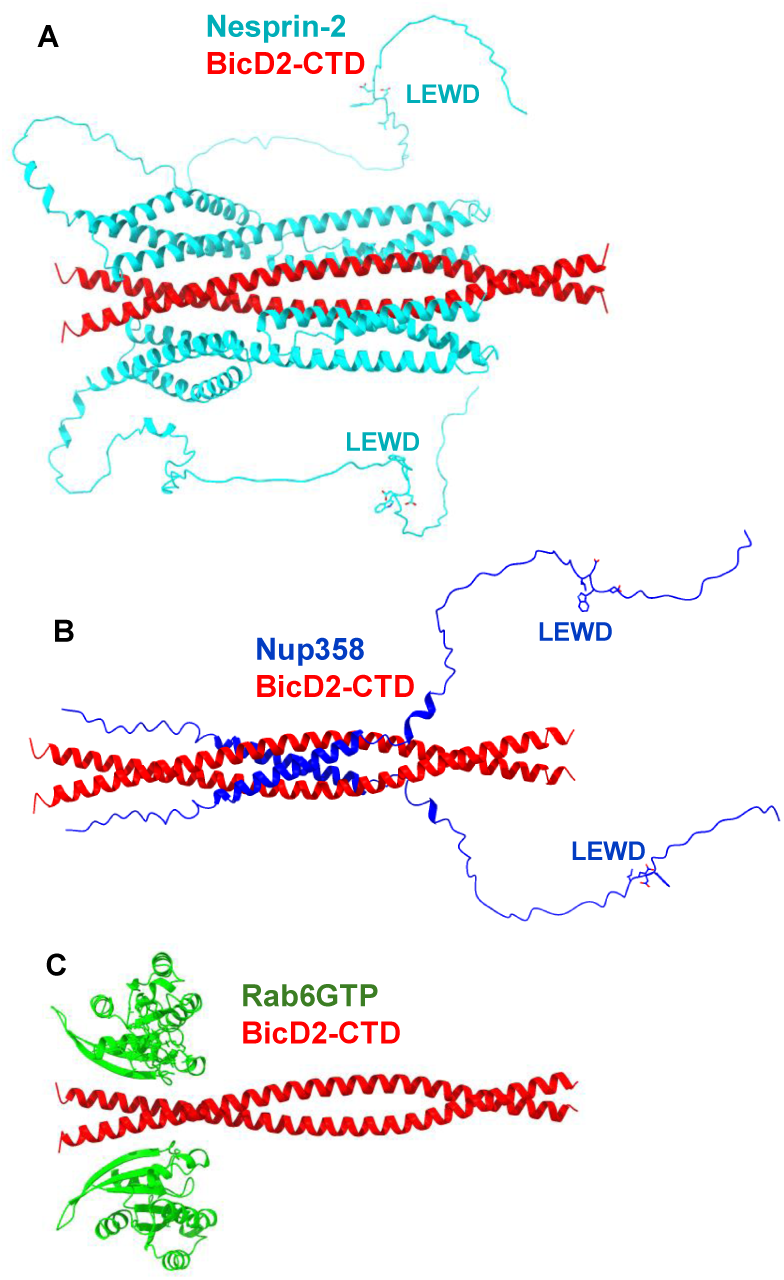
Cargoes bind to distinct but overlapping sites on BicD2-CTD. (A-C) The experimentally validated structural models of three minimal BicD2/cargo complexes are shown in cartoon representation, with BicD2-CTD in red ^24,34,36,47^. (A) Nesprin-2/BicD2 (Nesprin-2: cyan). (B) Nup358/BicD2 (Nup358: blue) ^34,36^. (C) Rab6^GTP^/BicD2 (Rab6^GTP^: green) ^24^. Note that Nup358 and Nesprin-2 have LEWD sequence motifs, which act as kinesin-1 binding site and are shown in stick representation. Note that in Nup358, the BicD2 binding site is separated from the LEWD motif by a ∼30 residue intrinsically disordered linker, while this linker consists of ∼65 disordered residues in Nesprin-2.

The minimal Nesprin-2/BicD2-CTD complex forms an alpha-helical bundle, in which SR52 and SR53 bind to BicD2-CTD. The BicD2 binding site is followed by a ∼65 residue intrinsically disordered domain and followed by a recruitment site for the kinesin-1 light chain 2 (Figure 5). Thus, the recruitment sites for dynein/BicD2 and kinesin-1 are spatially close but do not overlap, suggesting that the opposite polarity motors dynein and kinesin-1 may potentially be recruited simultaneously to Nesprin-2.

We carried out CD spectroscopy to assess if the alpha-helices of SR52 and SR53 are already present in apo-Nesprin-2 or whether they only fold upon binding to the BicD2-CTD (Figure S12). For these experiments, the purified minimal Nesprin-2 fragment and BicD2-CTD were mixed and incubated to assemble the complex, and CD wavelength scans were recorded. The CD wavelength scan of the complex is very similar to the sum of the CD wavelength spectra of the individual proteins (Figure S12). Both show comparable minima at 208 and 222 nm, which are characteristic for alpha-helical structures. The experimental error of the CD measurements is 3.5% at 208 nm and 5.2% at 222 nm ^36,59^. These results suggest that the minimal Nesprin-2/BicD2-CTD complex has a similar alpha-helical content compared to the individual proteins, suggesting that there are no large structural changes as observed for the Nup358/BicD2 complex ^36^.

The Nup358/BicD2 complex is structurally distinct (Figure S10, S11), but like the Nesprin-2/BicD2 complex, the BicD2 binding site of the Nup358/BicD2 complex is followed by a ∼30 residue intrinsically disordered domain and the LEWD motif (Figure 5) ^34,36,46^. The minimal BicD2 binding site on Nup358 is formed by a cargo-recognition alpha-helix, which binds to the center of the BicD2-CTD. This alpha-helix is intrinsically disordered in apo-Nup358 and folds to an alpha-helix upon binding to BicD2-CTD ^34,36^.

In the Rab6^GTP^/BicD2 complex, Rab6^GTP^ binds to the extreme C-terminal region of the BicD2-CTD (Figures 5, S10, S11), which is structurally distinct from the Nup358/BicD2 and Nesprin-2/BicD2 complex. In contrast to Nup358 and Nesprin-2, Rab6^GTP^ lacks a known kinesin-1 recruitment site. However, kinesin-1 in all three BicD2/cargo complexes is still recruited to the coiled-coil 2 domain of BicD2 ^25^, which binds via the kinesin-1 heavy chains. The interaction between BicD2 and cargoes is important to activate dynein for processive motility, and the observed structural differences of BicD2/cargo complexes (Figures S10, S11) could have implications for the overall motility of the associated dynein and kinesin-1 motors ^24,36^.

### Nesprin-2 fragments binds to full-length BicD2 and full-length kinesin-1

The interaction of full-length kinesin-1 hetero-tetramer (heavy and light chains) with three Nesprin-2 fragments was probed using single-molecule binding assays. The larger Nesprin-2 fragment (aa 6123-6421) that included the kinesin-1 recruiting LEWD motif (see Figure 2, Figure S13) showed a robust interaction with kinesin-1, with ∼20% dual-color images (Figure 6A, C). The other two Nesprin-2 fragments that did not include the LEWD motif showed only background levels of colocalization with kinesin-1 (∼5% for the Nesprin-2 aa 6123 – 6352 WT and 2%, for the V6246A/L6253A/F6261A triple mutant) (Figure 6C).

**Figure 6.**
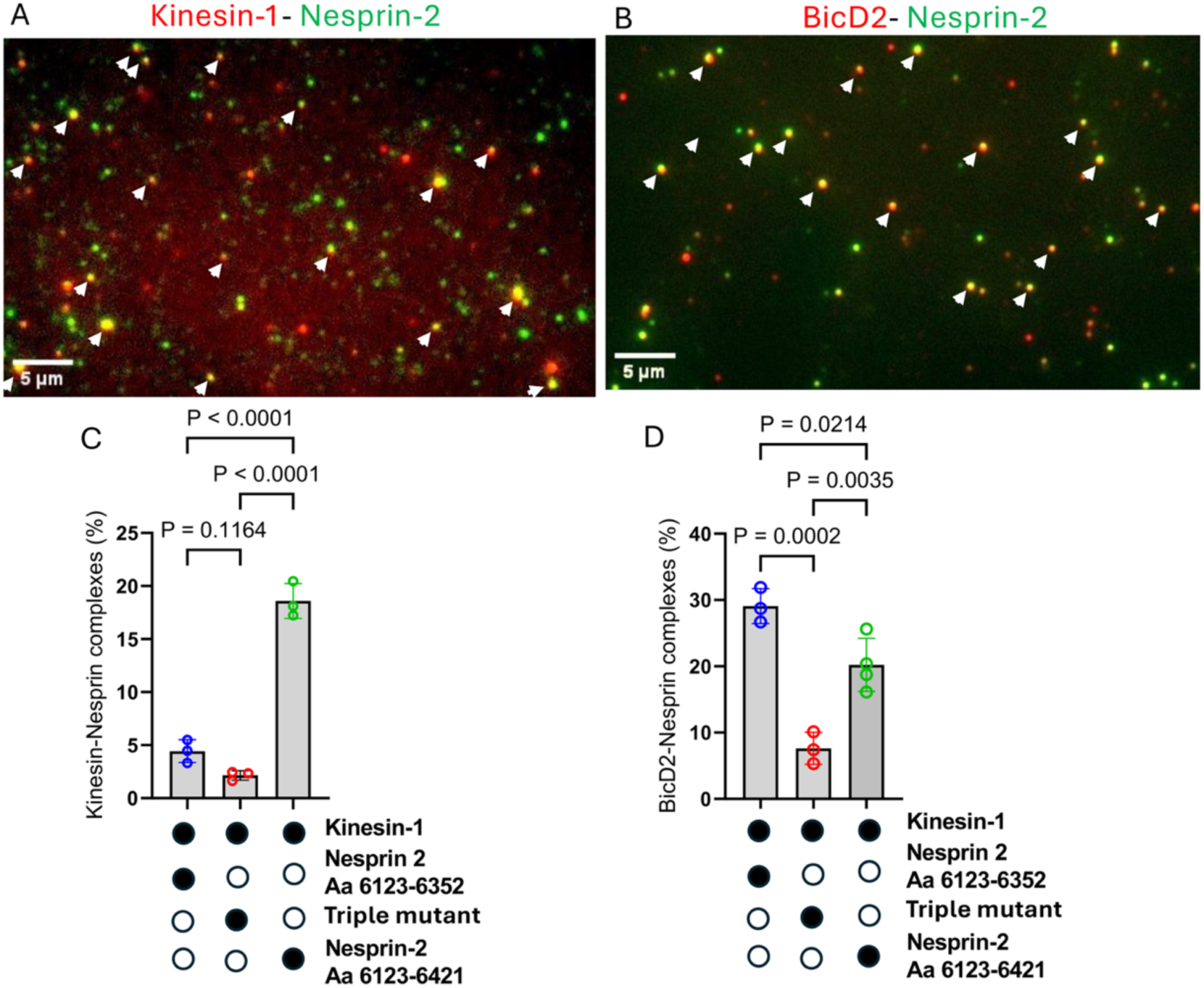
Minimal Nesprin-2 fragment interacts with full-length BicD2 and full-length kinesin-1. (A) Kinesin-1 (labeled with Alexa Fluor 647, red) and minimal Nesprin-2 fragment (aa 6123 - 6421, with GST-and SNAP-tag (for sequence see Figure S13), labeled with Alexa Fluor 488, green) interact, as indicated by yellow colocalized spots (white arrows). (B) Full-length BicD2 (labeled with Alexa Fluor 647, red) and minimal Nesprin-2 fragment (aa 6123 - 6352, with GST and Biotin-tag (Figure S13), labeled with Alexa Fluor 488, green) interact, as indicated by yellow colocalized spots (white arrows). (C) The quantification of the kinesin-1/Nesprin-2 co-localization is shown below for three Nesprin-2 fragments: 1) Nesprin-2 aa 6123-6352, which lacks the kinesin-1 recruiting LEWD motif; 2) the triple mutant (V6246A/L6253A/F6261A) of this fragment, which is expected to disrupt the interaction with BicD2. 3) Nesprin-2 aa 6123-6421, a larger fragment, which includes the LEWD motif. The filled circles below the graph indicate the components present in the assay. (D) Quantification of the BicD2/Nesprin-2 co-localization for the same three fragments described in C. The p values for (C, D) were calculated using one-way ANOVA followed by Tukey’s post-hoc test. Data were collected from three independent experiments.

We next probed the interaction of full-length BicD2 with the minimal Nesprin-2 fragment (aa 6123 – 6352) that interacted with BicD2 but lacked the LEWD motif that recruits kinesin-1 (see Figure 2). Full-length BicD2 and the minimal Nesprin-2 fragment showed 30% dual-colored images (Figure 6B, D), indicating a robust interaction. This co-localization was strongly diminished to ∼7% for the V6246A/L6253A/F6261A triple mutant of this fragment, confirming that this mutant also disrupts the Nesprin-2/BicD2 interaction in the context of full-length BicD2 (Figure 6D). We also assessed binding of BicD2 to a larger fragment of Nesprin-2 (aa 2123-6421) that included the LEWD motif (see Figure 2, this fragment was used for pulldown-assays and CD spectroscopy). This fragment showed a lower percentage (∼20%) of co-localization compared to the shorter Nesprin-2 (aa 6123 – 6352) fragment that lacked the LEWD motif (Figure 6D). A different labelling method (see Methods) was used for this fragment compared to the other ones (due to a SNAP tag versus a biotin tag), therefore the complex formation ratios are not directly comparable to the fragment without the LEWD motif. While the LEWD-motif containing domain of Nesprin-2 likely does not participate in the interaction with BicD2, it is possible that it has an allosteric effect on BicD2 binding, but this requires further investigation.

### One Nesprin-2 can activate the dynein/dynactin/BicD2 complex for processive motility

We evaluated the ability of the Nesprin-2 fragment that included the LEWD motif (aa 2123-6421) to relieve the autoinhibition of BicD2 and thus activate the BicD2/dynein/dynactin complex for processive motility. Dynein, dynactin, BicD2, and Nesprin-2 (DDBN) complexes were assembled at a molar ratio of 1:1:1:2 DDBN, with excess Nesprin-2 included to ensure binding to most BicD2 molecules. Single-molecule motility assays of these motor complexes on microtubules were performed using TIRF microscopy as previously described ^36,52^.

DDB complexes containing full-length auto-inhibited BicD2 or the constitutively active CC1 fragment of BicD2 (DDB^CC1^) were used as controls. Complexes were visualized by labeling full-length BicD2 or BicD2^CC1^ with Alexa 488 using a biotin–streptavidin conjugation system. The kymograph and movie for DDBN confirms activation for processive motion, while DDB lacking Nesprin-2 is mainly diffusive or static (Figure 7A and Movies S1, S2).

**Figure 7.**
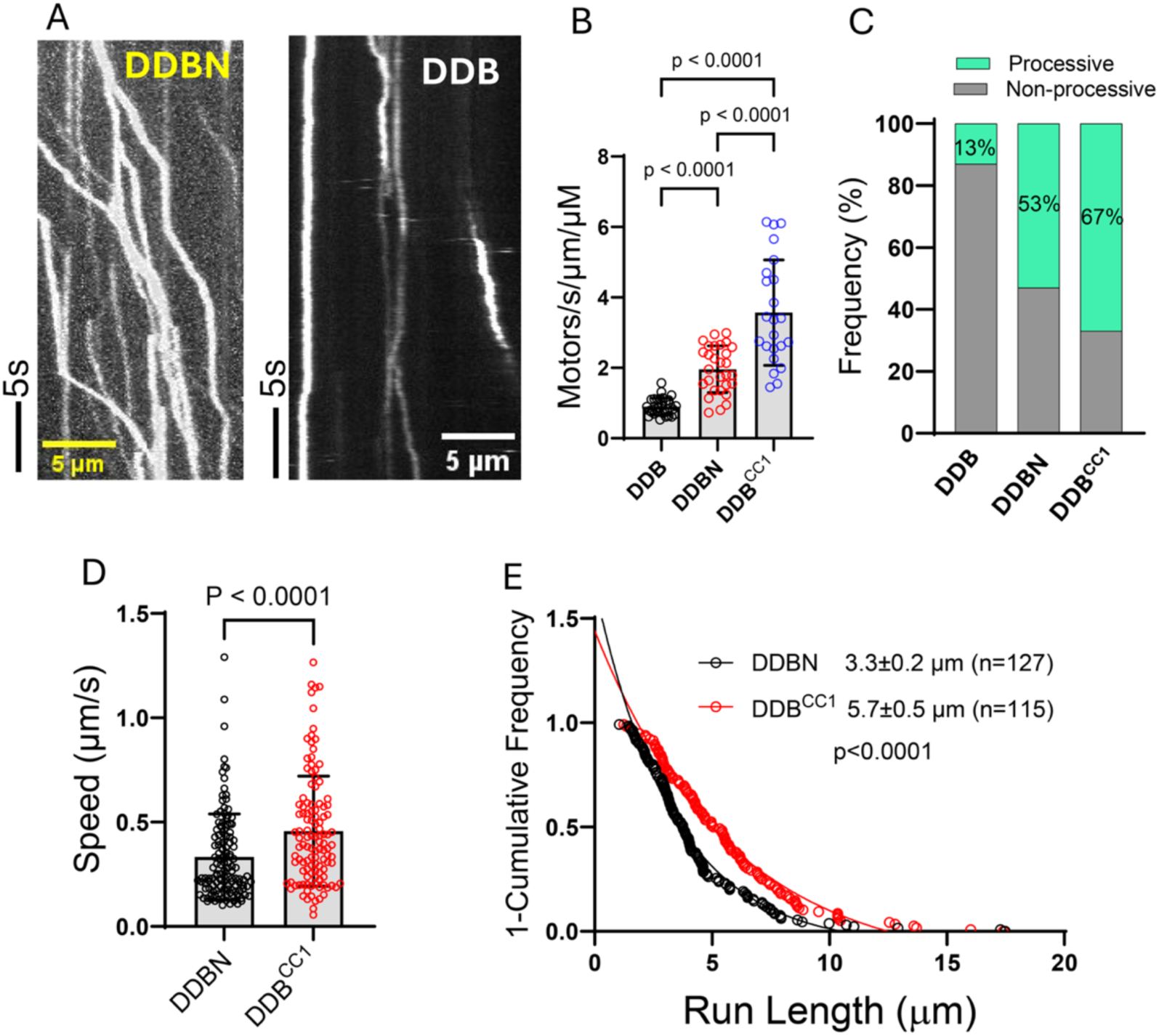
Nesprin-2 activates dynein/dynactin/BicD2 for processive motility. Single molecule processivity assays were performed to assess the motion of the assembled dynein/dynactin/BicD2/Nesprin-2 complex (DDBN) on microtubules. As controls, assays were also performed with autoinhibited dynein/dynactin + BicD2 (DDB) and the constitutively active dynein/dynactin/BicD2^CC1^(DDB^CC1^) complex. Labeled BicD2 alone showed negligible binding. (A) Kymographs for DDBN and DDB, showing that DDBN moves processively on MTs, while the DDB complex shows no or very little motion. See also Movie S1 (single-molecule processivity assay for DDB) and Movie S2 (single-molecule processivity assay for DDBN complexes). (B) Bar graph showing the binding rate (total number of microtubule-bound processive and static events) of DDBN per min per micrometer MT length, together with the DDB and DDB^CC1^ controls. The p values were calculated using one-way ANOVA followed by Tukey’s post-hoc test. (C) Bar graph showing the percent of the microtubule-bound motor complexes DDBN that were processive or non-processive. (D) The speed and (E) run length of DDBN was compared with the constitutively active DDB^CC1^ complex. The p-values were calculated using the Kolmogorov–Smirnov test. Data were collected from 3 independent experiments.

The total binding rate (processive and static microtubule-bound complexes combined) of DDBN to microtubules was 2.1-fold higher than for DDB (1.95 vs. 0.9 motors/s/µm/µM) (Figure 7B), but lower than that of the known active complex DDB^CC1^ ^36,50^.

Similarly, the percentage of moving DDBN complexes DDBN was 53%, more than three times greater than that of DDB (13%), but less than DDB^CC1^ levels (67%) (Figure 7C). The speed of DDBN (0.33 ± 0.20 µm/s, n=130) was slower than that of DDB^CC1^ (0.45 ± 0.26 µm/s, n=115) (Figure 7D), and the average run length of DDBN (3.3 ± 0.2 µm (n=127)) was shorter than DDB^CC1^ (5.7 ± 0.5 µm; n=115) (Figure 7E). The speed and run length of dynein/dynactin with autoinhibited full-length BicD2 (DDB) could not be determined due to the very low number of motile events, short travel distances, and frequent diffusion along microtubules. Taken together, the increased binding rate, high proportion of processive events, speed and run lengths indicate that Nesprin-2 relieves BicD2 autoinhibition, thereby activating the DDBN complex for processive motion. The level of activation remains lower than that observed for DDB^CC1^, but this is to be expected since the truncated BicD2 lacks the autoinhibition domain that is present in the complex with full-length BicD2 and Nesprin-2. Additionally, because BicD2 was the labeled species used for visualization, complexes without Nesprin-2 will remain in an auto-inhibited state and reduce the apparent level of activation.

This *in vitro* reconstitution of the Nesprin-2/BicD2/dynein/dynactin pathway with purified proteins confirms that no other components are required to activate this transport pathway. A previous study was performed with Nesprin-4, but it is structurally and functionally distinct from Nesprin-2 ^60^. The level of activation of dynein/dynactin/BicD2 by Nesprin-2 is similar to that seen with the alternative cargo adapter Nup358 ^36^.

To assess whether two Nesprin-2 molecules can bind to BicD2, and whether one or two molecules are required to activate the DDB motor complex, we reconstituted dynein/dynactin/BicD2/Nesprin-2 complexes with equal amounts of green and red labeled Nesprin-2 fragments and observed the complexes on microtubules (Figure 8A). While Nesprin-2 was labeled with either red or green Qdots, dynein, dynactin and BicD2 remained unlabeled. The yellow dots represent DDBN complexes containing two Nesprin-2 molecules bound, with 50% being the maximum statistically possible. Across three independent experiments, 11% of the complexes were dual colored yellow (n=509), corresponding to approximately 22% DDBN complexes with two bound Nesprin-2 molecules. In 78% of the complexes only one Nesprin-2 was bound. The observed percentage of dual-color complexes is a lower limit because various factors can reduce this number. As the labelling efficiency is not 100%, unlabeled Nesprin-2 molecules will lower the number of dual-color complexes. Additionally, the presence of a quantum dot on the first Nesprin-2 molecule may result in steric clashes that lower the affinity of the second Nesprin-2 molecule for BicD2. Separate experiments in which BicD2, instead of Nesprin-2, was labeled with two different colors showed that each DDBN complex contains only a single BicD (<2% dual colored, n=409), thus the yellow complexes observed when Nesprin-2 is labeled correspond to two molecules of Nesprin-2 bound to a single BicD2. We conclude that either one or two Nesprin-2 molecules can be recruited to DDBN complexes.

**Figure 8.**
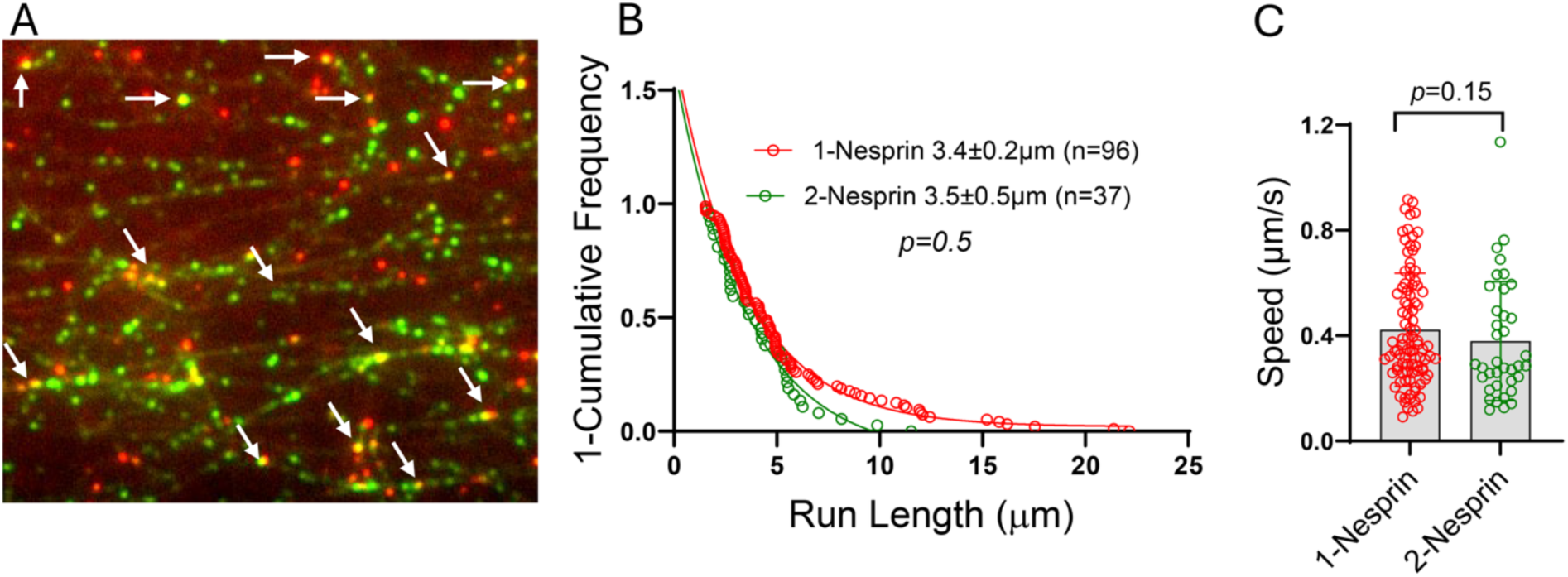
Binding of one or two Nesprin-2 fragments to BicD2 results in activation of DDBN complexes. (A) Single-molecule assay showing DDBN complexes, assembled using an equimolar mixture of red and green labeled Nesprin-2 fragments, bound to microtubules. Based on the number of single (red or green) and dual (yellow) colored images, 78% of DDBN complexes contain one bound Nesprin-2, while 22% of DDBN complexes contain two Nesprin-2 fragments. (B) Run lengths of DDBN complexes containing either one or two bound Nesprin-2 fragments are statistically the same, suggesting that one Nesprin-2 is sufficient to activate BicD2. (C) The speed of DDBN complexes is the same regardless of whether one Nesprin-2 or two Nesprin-2 fragments are bound. The mean speed of DDBN with 1-Nesprin bound was 0.42 ± 0.21 (n = 96), and with 2-Nesprin bound was 0.38 ± 0.22 (n = 37).

We next investigated whether one or two Nesprin-2 molecules are required to activate DDBN complexes for processive motion by comparing the run length and speed of DDBN complexes containing either one- or two Nesprin-2 molecules (Figure 8). For single-colored DDBN complexes, the run length was 3.4 ± 0.2 µm (n = 96), which is not statistically different from the run length observed when two Nesprin-2 molecules were bound (3.5 ± 0.5 µm, n = 37) (Figure 8B). Similarly, the velocities were statistically indistinguishable between the two conditions (0.42 ± 0.21 µm/s, n = 96 for single-colored vs. 0.38 ± 0.22 µm/s, n = 37 for dual-colored) (Figure 8C). These results suggest that a single Nesprin-2 molecule is sufficient to activate BicD2 for dynein motility. When we used AlphaFold to predict the structure of minimal Nesprin-2/BicD2 complexes with 1:2 and 2:2 stoichiometry (Figure 1), the predicted structural model of the 1:2 complex is similar to the structure of the 2:2 complex with only one Nesprin-2 binding site occupied, consistent with the result that either stoichiometry is capable of activating the DDBN complex for 2processive motion. In summary, single molecule experiments showed that Nesprin-2 binds kinesin via the LEWD motif, and binding of one molecule of Nesprin-2 to BicD2 is sufficient to activate the dynein/dynactin/BicD complex for motility.

## DISCUSSION

Here we present an experimentally validated structural model of the minimal Nesprin-2/BicD2 complex, where spectrin repeats (SR) 52 and SR53 of Nesprin-2 form an alpha-helical bundle with the BicD2 C-terminal cargo-binding domain (CTD). Either a single or two Nesprin-2 fragments are predicted to bind to symmetric binding sites formed by the BicD2 dimer. The intrinsically disordered domain, with the kinesin-1 recruiting LEWD motif is dispensable for the interaction with the BicD2-CTD, suggesting that both kinesin-1 and BicD2/dynein can potentially interact with Nesprin-2 simultaneously ^46^. We also show that the minimal Nesprin-2 fragment interacts with full-length BicD2 and activates BicD2/dynein/dynactin robustly for processive motility, suggesting that no additional components are required to activate this transport pathway. Interestingly, either a single or two Nesprin-2 molecules can be recruited to BicD2/dynein/dynactin complexes, resulting in robust activation of processive motility with similar speed and run length, supporting that both stoichiometries can be formed. Nesprin-2 bound DDBN complexes have the same speed and run length regardless of whether one or two Nesprin-2 are bound (Figure 8), suggesting that likely only one BicD2 dimer gets recruited to the motor complexes. It is surprising that a single Nesprin-2 fragment activates dynein/dynactin/BicD2 complexes, as formation of the BicD/Egalitarian complex requires a 2:2 stoichiometry to activate dynein/dynactin ^50^.

Overall, our results support 1:2 and 2:2 stoichiometries for Nesprin-2/BicD2 complexes, dependent on the context (i.e. the stoichiometry may be affected by tags, concentration, membrane tethering or the Nesprin-2 oligomerization state). The oligomeric state of the cytoplasmic Nesprin-2 domain is matter of debate. It has been established that the KASH/SUN interaction in the lumen of the nuclear envelope is formed by three Nesprin-2 and three SUN-proteins ^61^. A recent study provided evidence for regulation of the oligomeric state of the cytoplasmic Nesprin-2 domain. In the absence of F-actin, the domain formed monomers, but the interaction with F-actin stimulated oligomerization ^5^. It is conceivable that a similar activation of BicD2 by either a single or two Nesprin-2 molecules as observed here may serve to regulate dynein motility in a manner that is independent from the oligomeric state of Nesprin-2 in cells, which may change upon recruitment of F-actin ^5^. It should also be noted that Nesprin-2 is expressed as distinct isoforms with distinct domain compositions.

A recent paper reported that additional regions of Nesprin-2, notably SR48, markedly increased the affinity of BicD2 to Nesprin-2 ^40^, which is not present in our minimal Nesprin-2 fragment. SR48 is however not essential for motor recruitment and healthy brain development: In the mini-Nesprin-2, the N-terminal actin-recruiting domain is fused to the microtubule motor recruitment domain located in SR52 - SR56, followed by the transmembrane alpha-helix and the C-terminal KASH domain. It should be emphasized that while depletion of Nesprin-2 causes defects in neuronal migration in rat embryo brains, expression of a mini-Nesprin-2 that included the microtubule motor recruitment domain (SR52-SR56) but lacked SR48 restored near normal migrations of neurons in the cortical plate, suggesting that SR48 is dispensable for the recruitment of kinesin-1 and dynein to Nesprin-2 ^1,2^. Furthermore, we show that the minimal Nesprin-2 fragment composed of SR52 and SR53 interacts with full-length BicD2 and activates BicD2/dynein/dynactin complexes robustly for processive motility. According to established models of BicD2 activation, which may also apply to activation by Nesprin-2, full-length BicD2 forms a looped, auto-inhibited conformation that binds dynein–dynactin weakly and does not result in activation, as the N-terminal dynein binding site is sterically occluded by the C-terminal cargo binding domain (CTD) ^50,62,63^. The CTD of BicD2 is required for auto-inhibition, as a truncated BicD2 fragment without this domain (BicD2^CC1^) is constitutively active. Binding of cargo to the CTD results in a conformational change that promotes loop-opening and activates full-length BicD2 for dynein and dynactin recruitment, and cargo-loaded adapters such as BicD2 are required for activation of processive motility ^63,64^. We have shown previously that specific interactions between Nup358/BicD2 and Rab6/BicD2 are important for overall motility of the recruited dynein complexes. Differences in speed and run length were observed for Nup358 mutants that disrupted the interaction with BicD2 ^24,36^, suggesting that structurally distinct BicD2/cargo interactions may modulate overall motility in cellular transport pathways.

The structural model of the minimal Nesprin-2/BicD2 complex is structurally distinct from other BicD2/cargo complexes but shares the LEWD motif with Nup358. As with Nup358, the Nesprin-2 LEWD motif is located in an intrinsically disordered domain C-terminal of the BicD2 binding site. We show that this disordered domain with the LEWD motif is dispensable for BicD2 binding, whereas a recent study indicated that the mutation of the LEWD motif of Nesprin-2 to LEAA diminished binding to BicD2 in a pulldown assay that included cell extracts ^2^. The LEAA mutation also impacted neuronal migration with fewer neurons completing it and reaching the cortical plate ^2^. While our experiments (Figures 1 and 2) suggest that the LEWD containing domain of Nesprin-2 is dispensable for BicD2 binding, future studies are required to investigate if this domain may modulate binding of BicD2 though allosteric effects that may also potentially involve kinesin-1. The intrinsically disordered linker between the BicD2 binding site and the LEWD motif is substantially longer for Nesprin-2/BicD2 compared with Nup358/BicD2 (∼65 versus ∼30 residues) which may impact the overall coordination of dynein and kinesin-1, resulting in distinct motilities. In contrast, Rab6 lacks a LEWD motif or another known kinesin-1 recruiting site. In all three BicD2/cargo complexes, the kinesin-1 heavy chain is also recruited via the coiled-coil 2 domain of BicD2 ^25^. The observed structural differences of the BicD2/cargo interactions as well as the observed distinctions in the adjacent kinesin-1 recruiting sites could finetune overall motility of the recruited dynein and kinesin-1 complexes.

How BicD2’s preference for these three distinct transport pathways (Rab6, Nesprin-2 and Nup358) during brain development and other physiological processes remains to be established. One of the key regulators of the Nup358 pathway is the G2 phase specific kinase cyclin dependent kinase 1 (Cdk1), which phosphorylates Nup358 and increases its affinity towards BicD2 ^65^. At the same time, Cdk1 phosphorylates Rab6 with the effect of lowering its affinity to BicD2 ^66^. Furthermore, BicD2 is also phosphorylated by Cdk1 at residue Ser102, which promotes recruitment of PLK1 (polo-like kinase 1) and increases its affinity for Nup358. It is conceivable that the Nesprin-2/BicD2 nuclear positioning pathway is regulated by kinases as well. Thirteen phosphorylation sites were identified in the minimal domain of Nesprin-2, which bind to BicD2 and kinesin-1 (aa 6123-6421; Table S3), and 10 phosphorylation sites were identified in the same domain of Nesprin-1 (Table S4), but their effect on the interaction with BicD2 remains to be established. Interestingly, residue T8033 of Nesprin-1, which is located in the BicD2 binding domain and corresponds to T6157 in mouse Nesprin-2 (Figure S4), is phosphorylated by PLK1 (Table S4), which is recruited in G2 phase to BicD2 ^67^. Future studies will establish the role of this phosphorylation site on regulation of this pathway. It is conceivable that the phosphorylation by PLK1 could either diminish or increase recruitment of Nesprin-1/2 to BicD2 and specifically down-or up- regulate this nuclear positioning pathway during interkinetic nuclear migration towards the apical brain surface, which occurs in G2 phase, when PLK1 is active.

Several disease mutations that cause brain and muscle developmental diseases in infants, including spinal muscular atrophy, are located in the cargo binding domain of BicD2 ^18,32,68,69^. We have previously shown that several of the disease mutations affect the affinity of BicD2 towards Nesprin-2, Nup358 and Rab6 in a distinct manner, thereby modulating associated cellular transport pathways that are important for brain development ^18^. For example, the R747C and E774G mutations which cause spinal muscular atrophy decrease binding to Nup358 but increase binding to Nesprin-2 ^18^. This is correlated with defects in the Nup358-dependent nuclear positioning pathway that is essential for apical nuclear migration, resulting in brain development defects ^18^. Interestingly, the R747C mutation does not affect the interaction with Rab6, while the E774G mutation diminishes binding to Rab6 ^32,59,62^. The BicD2 R694C mutation on the other hand, which causes arthrogryposis multiplex congenita with polymicrogyria, which may result in infants death ^70^, has a 3-fold higher affinity towards Nup358. This mutation results in less recruitment of Nesprin-2 to BicD2 and is associated with defects in the Nesprin-2 associated nuclear positioning pathway that is important for neuronal migration ^18^. It should be noted that in addition to the BicD2 R694C mutation, a Nesprin-1 mutation has been identified that also causes arthrogryposis ^71^. Another mutation that reduces Nesprin-2 binding is the BicD2 K774Ter truncation variant which causes lissencephaly ^20,72^. A recent publication also noted extensive proteome changes including gain of function interactions for several BicD2 disease mutations ^73^. These results are in line with our structural models of three BicD2/cargo complexes, according to which Nesprin-2, Nup358 and Rab6 bind to overlapping but distinct sites on BicD2 ^24,59^ (Figure 5, Figure S10, S11).

Mutations in Nesprin-1 and Nesprin-2 also cause Emery-Dreifuss muscular dystrophy (EDMD) ^41^. Interestingly, four human mutations that cause EDMD were identified in the minimal Nesprin-1 and Nesprin-2 domains that bind to BicD2 and kinesin-1. The Nesprin-1 R8095H and R8212H mutations as well as the T6211M mutation of Nesprin-2 are all located in the spectrin repeats that act as BicD2 binding site (Figures S4 and S5) ^41,2^. The EDMD causing R8272Q mutation of Nesprin-1 is located close to the LEWD motif and diminishes the affinity to the kinesin LC2 ^41^. The mutation also reduced recruitment of kinesin-1 to the nuclear envelope in myotubes and resulted in reduced myonuclear fusion ^41^. Interestingly, depletion of kinesin LC2 in myotubes caused a similar phenotype ^41^. It is conceivable that all four mutants may affect dynein and kinesin-1 mediated myonuclear positioning during muscle cell differentiation, thereby contributing to EDMD pathogenesis. Of note, abnormally clustered nuclei have been found in patients with EDMD, suggesting that correct nuclear positioning may be required for proper muscle functions ^15^.

## CONCLUSIONS

A structural model of the minimal Nesprin-2/BicD2 complex was established and experimentally validated. Either a single or two Nesprin-2 fragments are predicted to bind to symmetric binding sites on the BicD2 dimer. The core of the interaction is formed by spectrin repeats of Nesprin-2 that form a helical bundle with BicD2, which is structurally distinct from other BicD2/cargo complexes. The structurally distinct interactions may modulate overall motility of the associated transport pathways. The BicD2/dynein binding site on Nesprin-2 is spatially close but does not overlap with the kinesin-1 binding site. We have established that recruitment of either one or two Nesprin-2 fragments to the BicD2/dynein/dynactin complex activates it robustly for processive motility, resulting in similar speed and run length. This suggests that no other components are needed for this transport pathway and paves the way for future mechanistic studies. We have also designed a triple mutant that disrupts the interaction between BicD2 and Nesprin-2, which will serve as a valuable tool to study the role of this nuclear positioning pathway in brain development. Several EDMD causing mutations of Nesprin-1 and 2 are located in the motor recruitment domain and may alter interactions with BicD2 and kinesin-1, in line with the observation of abnormally clustered nuclei in patients with EDMD. Our results will guide future studies of the essential roles of Nesprin-1/2 in brain and muscle development and reveal underlying causes for devastating brain and muscle developmental diseases caused by Nesprin-1/2 and BicD2 mutations including EDMD and spinal muscular atrophy.

## SUPPORTING INFORMATION

The supporting information (PDF) contains 4 supporting tables (Nesprin-2/BicD2 contact residues and Nesprin-2 phosphorylation sites), 13 supporting figures (SEC-MALS data, sequence alignments, AlphaFold structure predictions, control experiments for Figure 3, structural analysis of BicD2/cargo complexes, and CD data), supporting methods (SEC-MALS) and supporting references, as well as two supporting movies with single-molecule processivity assays (AVI).

## ACKNOWLEDGEMENTS

SR Solmaz and MY Ali were supported by NIH grant R01 GM144578. The project was also supported by NIH grant R35 GM136288 to KM Trybus and NIH grant R03 NS126811-01A1 to MY Ali. The CD instrument was supported by NIGMS grants 1R01GM125853-02S1 and 3R35GM130207-01S1. The authors declare that they have no conflict of interest.

## TABLE OF CONTENT FIGURE

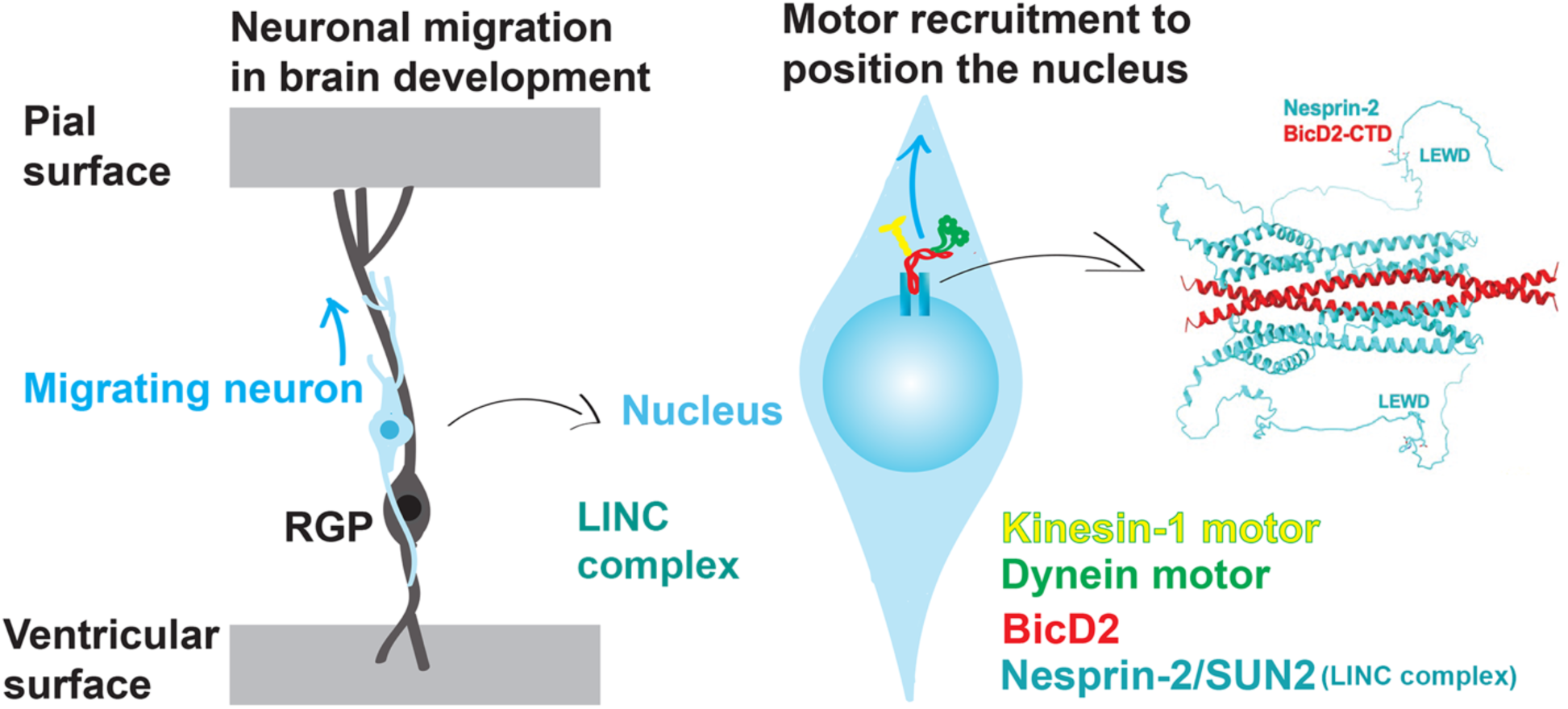

