## Supplemental Tables and Figures for "Cargo recognition of Nesprin-2 by the dynein adapter Bicaudal D2 for a nuclear positioning pathway that is important for brain development"

**Table S1. Summary of contact residues from PISA. Experimentally validated contact residues are denoted by an asterisk.**

| **Nesprin-2/BicD2**<br>**Nesprin-2**<br>**aa 6123-6421** | **Nesprin-2/BicD2**<br>**BicD2**<br>**aa 715-804** | **Rab6/BicD2**<br>**BicD2**<br>**aa 715-804** | **Nup358/BicD2**<br>**BicD2**<br>**aa 715-804** |
|---|---|---|---|
| 6124 | | | 732 |
| 6160 | 734 | | |
| 6164 | | | 735 |
| 6171 | | | 736 |
| 6174 | | | 737 |
| 6175 | 738 | | |
| 6178 | | | 739 |
| 6181 | | | 740 |
| 6182 | 741 | | 741 |
| 6185 | 742 | | 742 |
| 6189 | | | 743 |
| 6193 | | | 744 |
| 6195 | 745 | | |
| 6241 | 746 | | 746 |
| 6242 | | | 747 |
| 6245 | 749 | | 749 |
| 6246* | | | 750 |
| 6249 | | | 751 |
| 6250* | 753 | | 753 |
| 6252 | | | 754 |
| 6253* | | | 755 |
| 6254 | 756 | | 756 |
| 6256 | 757 | | 757 |
| 6257 | | | 758 |
| 6259 | | | 759 |
| 6260* | 760 | | 760 |
| 6261* | | | 761 |
| 6312 | | | 762 |
| 6315 | | | 763 |
| 6319* | | | 764 |
| 6322 | 765 | | 765 |
| 6323 | | | 766 |
| 6326 | | | 767 |
| 6329* | | | 768 |
| 6333 | | | 769 |
| 6337 | | | 770 |
| | 772 | | 772 |
| | 774 | 774 | 774 |
| | 775 | | 775 |
| | | | 776 |
| | 777 | 777 | |
| | 778 | 778 | |
| | 779 | 779 | 779 |
| | 781 | 781 | 781 |
| | | 782 | 782 |
| | 783* | 783 | 783 |
| | 784 | 784 | |
| | 785 | 785 | 785 |
| | 786* | 786 | 786 |
| | 787 | 787 | |
| | 788* | 788 | 788 |
| | 789* | 789 | 789 |
| | 790 | 790 | 790 |
| | 791 | | |
| | 792 | 792 | |
| | 793 | 793 | 793 |
| | 794 | 794 | |
| | 795 | 795 | 795 |
| | 796 | | |
| | 797* | 797 | 797 |
| | | 798 | |
| | 800 | | |
| | 801 | 801 | 801 |
| | 804 | 804 | |

See also Figure 3.

**Table S2. Secondary structure estimation from the CD spectra with BeStSel** [1]

| Nesprin-2 mutant | α-helix (%) | Anti-parallel β-sheet (%) | Parallel β-sheet (%) | Turn (%) | Other (%) |
|---|---|---|---|---|---|
| **WT** | 51.9 | 0 | 0 | 11.2 | 36.9 |
| **V5695A** | 45.6 | 1.8 | 1.9 | 12.0 | 38.6 |
| **E6250A** | 48.4 | 8.3 | 0.1 | 11.3 | 31.9 |
| **H6260A** | 48.8 | 0 | 0 | 11.1 | 40.1 |
| **V6246A/L6253A/F6261A** | 52.7 | 1.8 | 0 | 9.7 | 35.8 |
| **R6329A** | 52.2 | 15.6 | 0 | 19.4 | 12.7 |
| **Y6319A** | 48.1 | 2.2 | 0 | 9 | 40.8 |

**Table S3. Nesprin-2 phosphorylation sites from the PhosphoSitePlus v 6.8.1 database**

| Phosphorylation Site (human) | Phosphorylation Site (mouse) | References |
|---|---|---|
| S6314 | S6301 | [2] |
| T6348 | T6335 | [3] |
| S6349 | S6336 | [3–5] |
| T6351 | | [5] |
| S6361 | S6348 | [6] |
| T6365 | T6352 | [7] |
| | | [8] |
| S6384 | S6371 | [2] |
| | T6375 | [9] |
| S6389 | S6376 | [8] |
| S6392 | S6379 | [9,10] |
| | S6400 | [9] |
| | S6403 | [9] |

Note that the minimal BicD2 binding site was mapped to aa 6123 - 6352 of mouse Nesprin-2, which is followed by the adaptive domain (aa 6353 - 6421).

**Table S4. Nesprin-1 phosphorylation sites.**

**Nesprin-1 phosphorylation sites with known kinase specificity from the PhosphoSitePlus v 6.8.1 database**

| Phosphorylation Site (human) | Phosphorylation Site (mouse) | Kinase | Reference |
|---|---|---|---|
| T8033 | | **PLK1** | [11–13] |

**Nesprin-1 phosphorylation sites from the PhosphoSitePlus v 6.8.1 database**

| Phosphorylation Site (human) | Phosphorylation Site (mouse) | Source |
|---|---|---|
| T8033 | | [11–13] |
| S8223 | S8227 | [14] |
| S8232 | | [15] |
| S8236 | S8240 | [15] |
| S8250 | S8254 | [5] |
| S8270 | | [16] |
| S8274 | S8278 | [9,16,17] |
| S8277 | S8281 | [18] |
| S8280 | S8284 | [18] |
| S8305 | S8308 | [6] |

Note that the BicD2 recruiting SR70 – SR71 are located in aa 7998 -8216 of mouse Nesprin-1, which is followed by the adaptive domain.

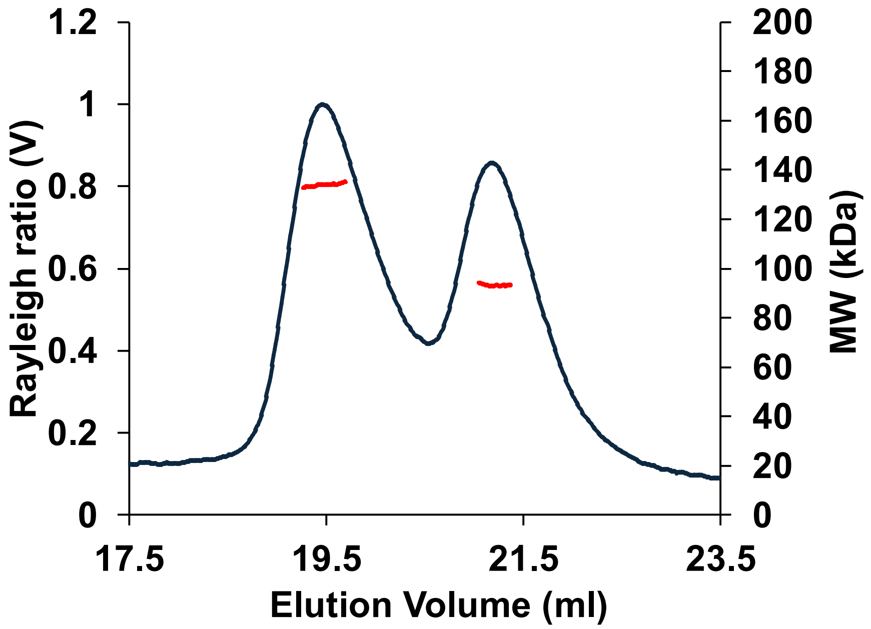


**Figure S1. Size exclusion chromatography coupled to multi-angle light scattering (SEC-MALS) data suggest a 2:2 stoichiometry for the minimal Nesprin-2/BicD2 complex.** 0.225 mg/ml Nesprin-2 (GST-tagged, aa 6123-6352; for sequence see Figure S13) and 0.04 mg/ml BicD2-CTD (aa 715-804, the $his_6$-tag was proteolytically cleaved) were mixed and incubated for 30 min prior to SEC-MALS analysis. The Rayleigh ratio (blue) and weight-averaged molar masses (MW in kilodaltons, red) are plotted versus the elution volume (ml) for a representative experiment. Molar masses were determined across the selected peak area and plotted on a secondary axis. The average MW from four experiments is MW=128.9 ± 4.9 kDa for the left peak, which closely matches the calculated mass of a 2:2 complex of Nesprin-2 and BicD2 (130.4 kDa; note that the mass of a BicD2 protomer is 10.9 kDa, and the mass of a Nesprin-2 protomer is 54.3 kDa). For the right peak, MW= 94.1 ± 2.2 kDa. The standard deviation was calculated as error.

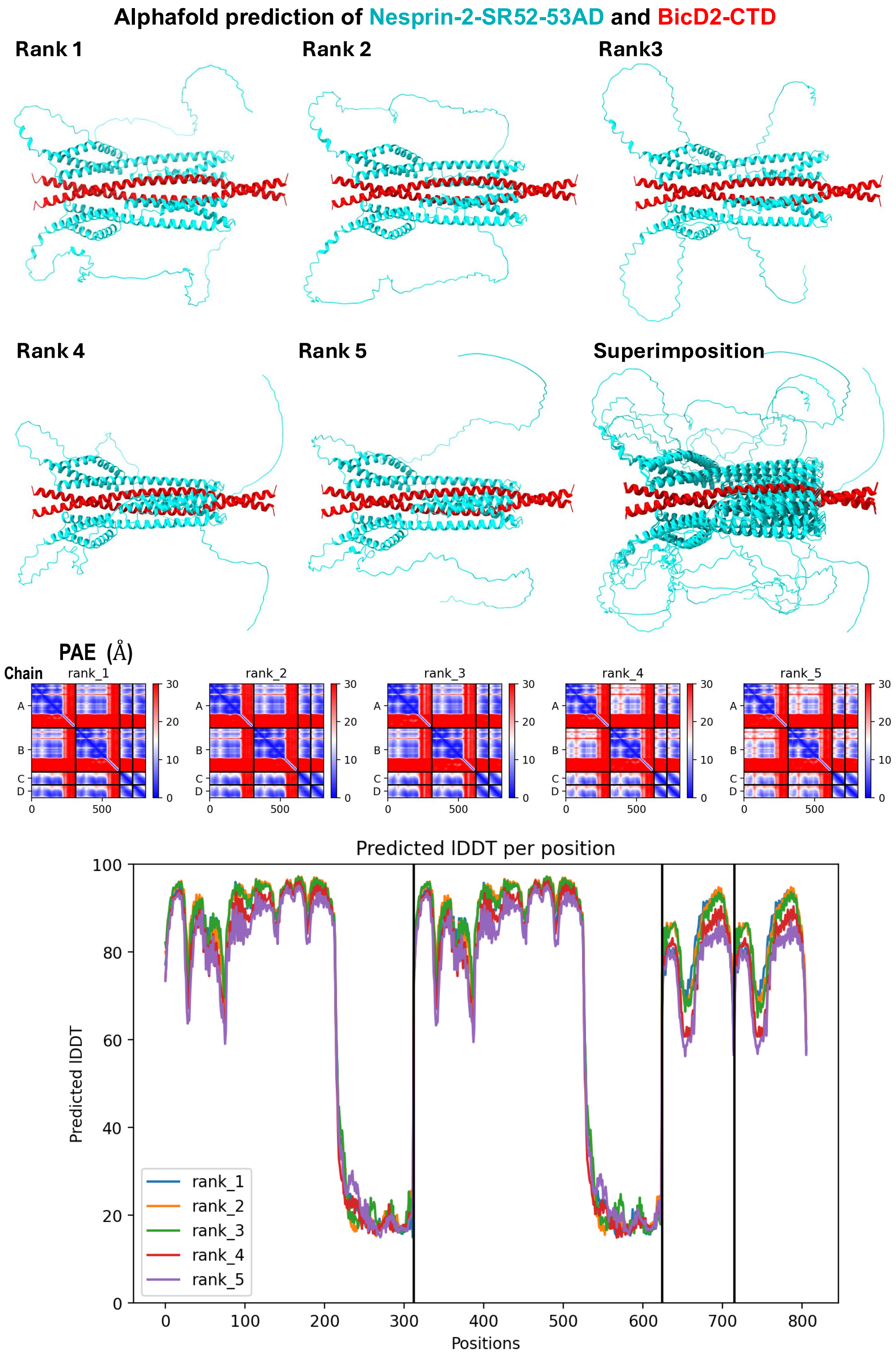


**Figure S2. AlphaFold prediction of the minimal Nesprin-2/BicD2** complex (Nesprin-2-SR52-SR53AD and BicD2-CTD). The five highest ranked predicted structures are shown individually and superimposed in cartoon representation, together with the PAE and pLDDT error plots. See also Figure 1.

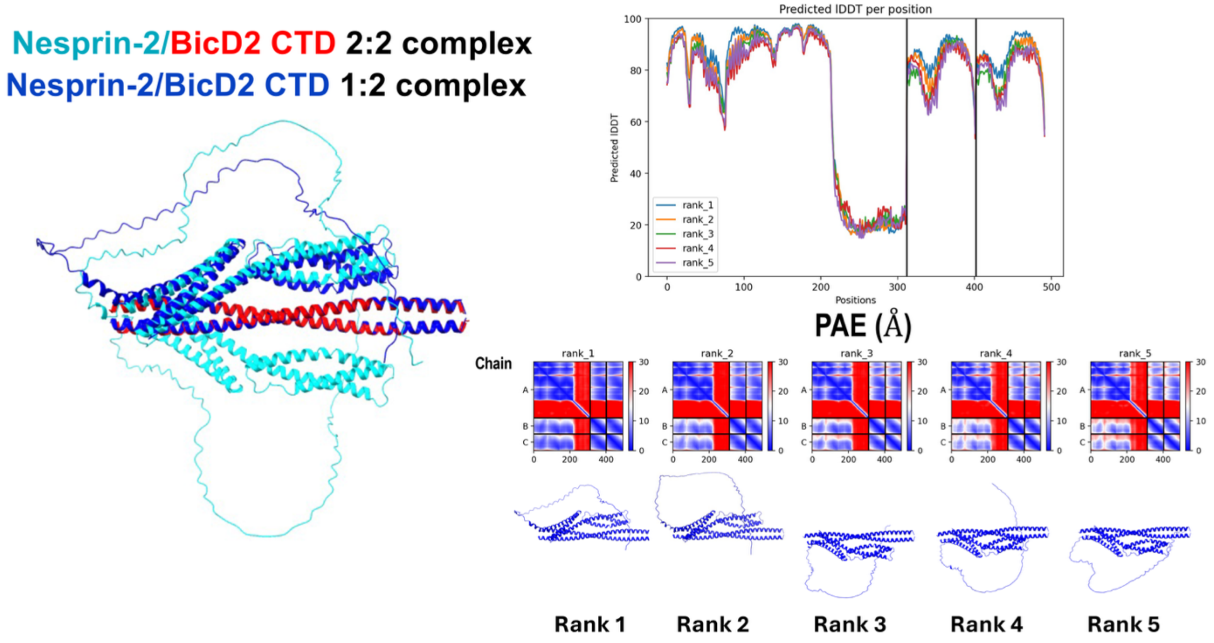


**Figure S3. AlphaFold prediction of the minimal Nesprin-2/BicD2 complex with a single Nesprin-2 molecule bound to two BicD2 molecules in blue** (Nesprin-2-SR52-SR53AD and BicD2-CTD). The prediction is least-squares superimposed with the 2:2 complex of Nesprin-2 and BicD2 in cyan and red. The RMSD is 1.8 Å, which suggests that the predicted binding mode of a single Nesprin-1 is very similar to the binding mode of two Nesprin-2 molecules (note that the disordered regions have the highest RMSD). The cartoon representations of the five highest-ranking models are shown along with their PAE and pLDDT error plots. See also Figure 1.

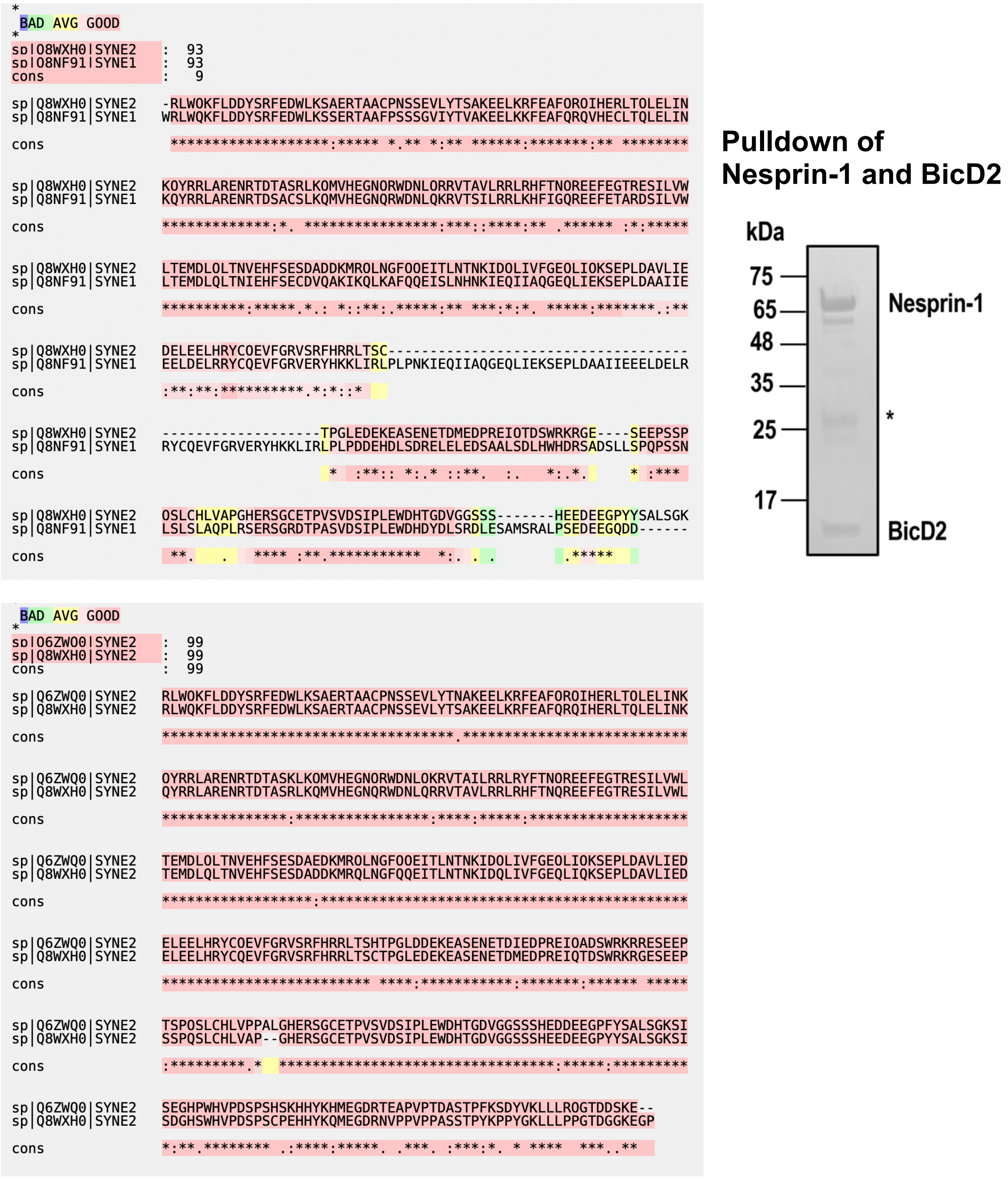


**Figure S4. Top: A sequence alignment of human Nesprin-2-SR52-SR53AD and human Nesprin-1-SR70-SR71AD confirms high sequence conservation** (prepared with T-COFFEE, version 11.00). SDS PAGE analysis of the elution fraction from a pull-down assay confirms that the GST-tagged Nesprin-1-SR70-SR71AD-GST fragment interacts with the His$_6$-tagged BicD2-CTD. The asterisk denotes GST. Masses of standard proteins are indicated**.** Bottom: **sequence alignment of mouse and human Nesprin-2-SR52-SR53AD**. Uniprot accession numbers for the sequences are shown. The sequence conservation (cons) is illustrated by a color gradient from red (good) to blue (bad), for which the scale is shown at the top, together with the overall sequence conservation. Asterisks denote identical residues, colons denote highly conserved residues, full stops indicate partially conserved regions and blank spaces denote residues that are not conserved. See also Figure 1.

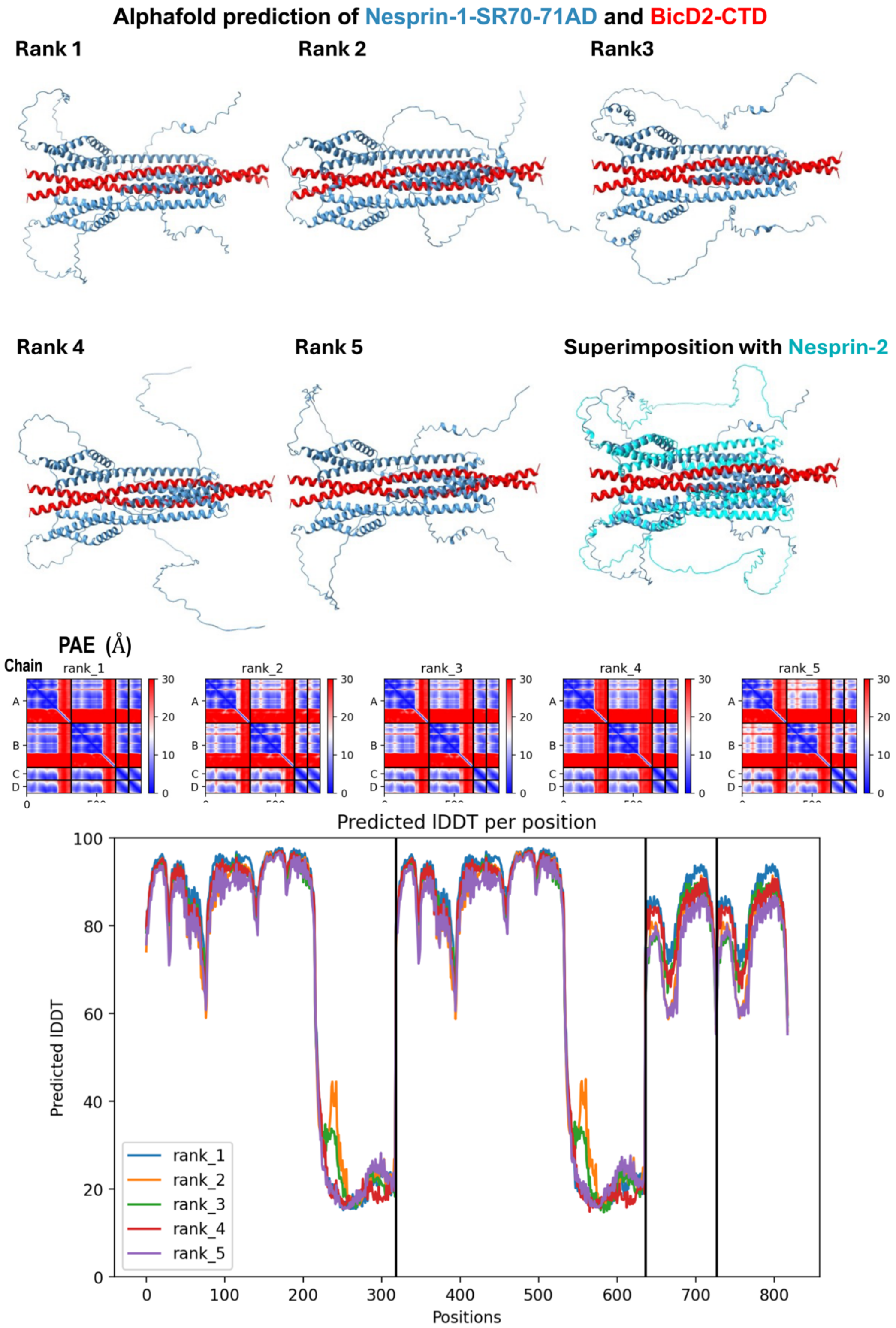


**Figure S5. AlphaFold prediction of the Nesprin-1-SR70-SR71AD and BicD2-CTD complex.** The five highest ranked predicted structures are shown in cartoon representation, together with the PAE and pLDDT error plots. See also Figure 1.

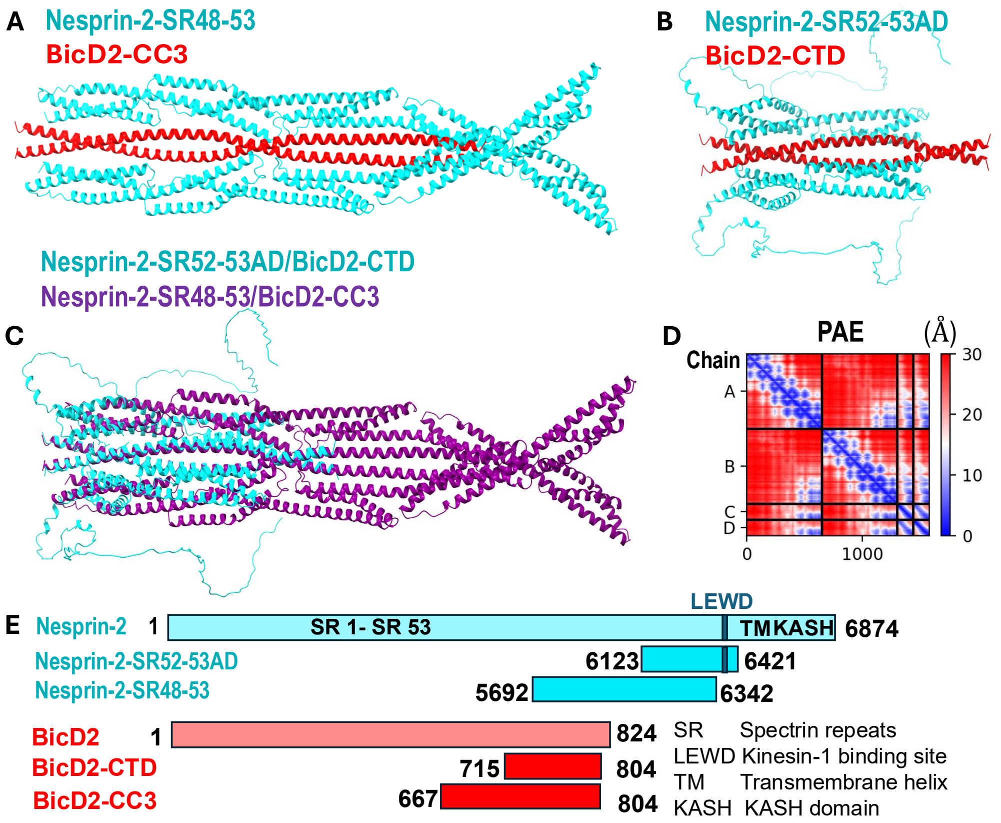


**Figure. S6. The predicted structure of a larger Nesprin-2/BicD2 complex including SR 48- SR 58 and the entire coiled coil 3 of BicD2 supports the predicted structure of the minimal complex.** (A) Structural model of a larger Nesprin-2/BicD2 complex (Nesprin-2 aa 5692-6342 [19] and BicD2 aa 667-804) from AlphaFold, shown in cartoon representation. (B) Structural model of the minimal complex (Nesprin-2 aa 6123-6421 and BicD2 aa 715-804). (C) Least squares superimposition of A (purple) and B (cyan) (RMSD = 0.99 Å). Note that the predicted structure of the minimal complex closely resembles the predicted structure of the same domains in the context of the larger complex. (D) Predicted aligned error (PAE) plot for the highest-ranking model. The x and y axes show the residue numbers, starting with Nesprin-2 (Chain A, B), followed by BicD2 (Chain C, D). Note that the high PAE suggests that the predicted structure of the additional regions in the larger complex is not reliable. Additional models and error plots are shown in Figure S7. (E) Schematic representation of the Nesprin-2 and BicD2 fragments used for A and B [19–21]. See also Figure 1.

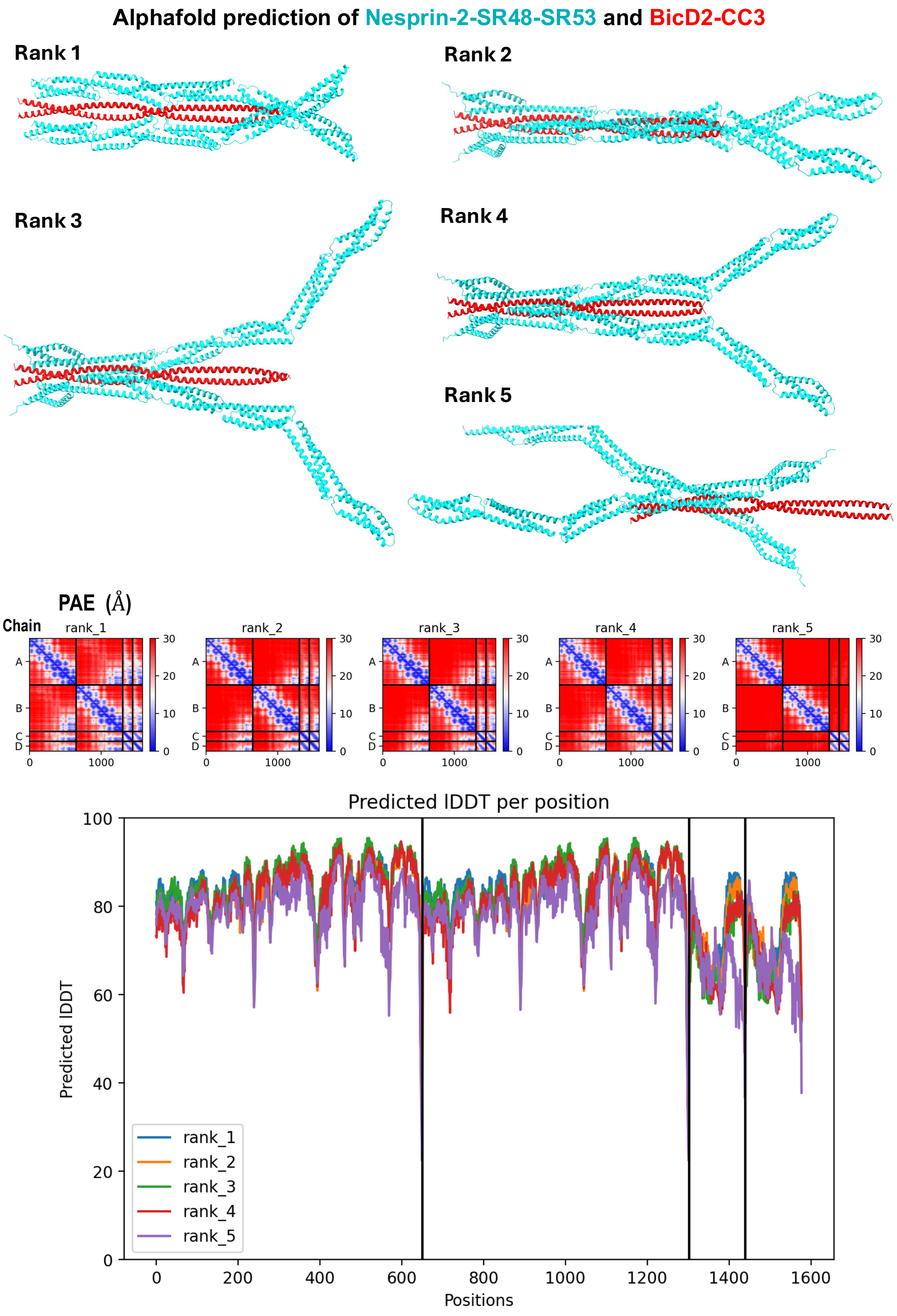


**Figure S7. AlphaFold prediction of the Nesprin-2-SR48-SR53 and BicD2-CC3 complex.** The five highest ranked predicted structures are shown in cartoon representation, together with the PAE and pLDDT error plots. Note that the PAE and pLDDT error suggest that the structure prediction of the additional regions in the larger complex is not reliable. See also Figure S6

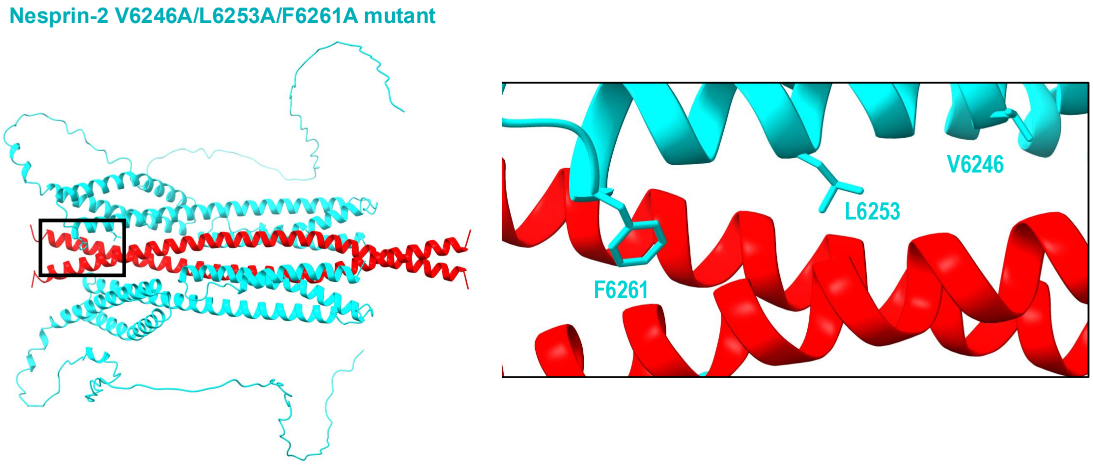


**Figure S8. The Nesprin-2/V6246A/L6253A/F6261A triple mutant virtually abolishes the interaction with the BicD2-CTD.** Cartoon representation of the minimal Nesprin-2/BicD2 complex. The boxed region on the left is enlarged on the right to depict the location of the mutated residues. Residues F6261, L6253 and V6246, which mediate contacts with BicD2 are shown in stick representation and labelled. See also Figure 3.

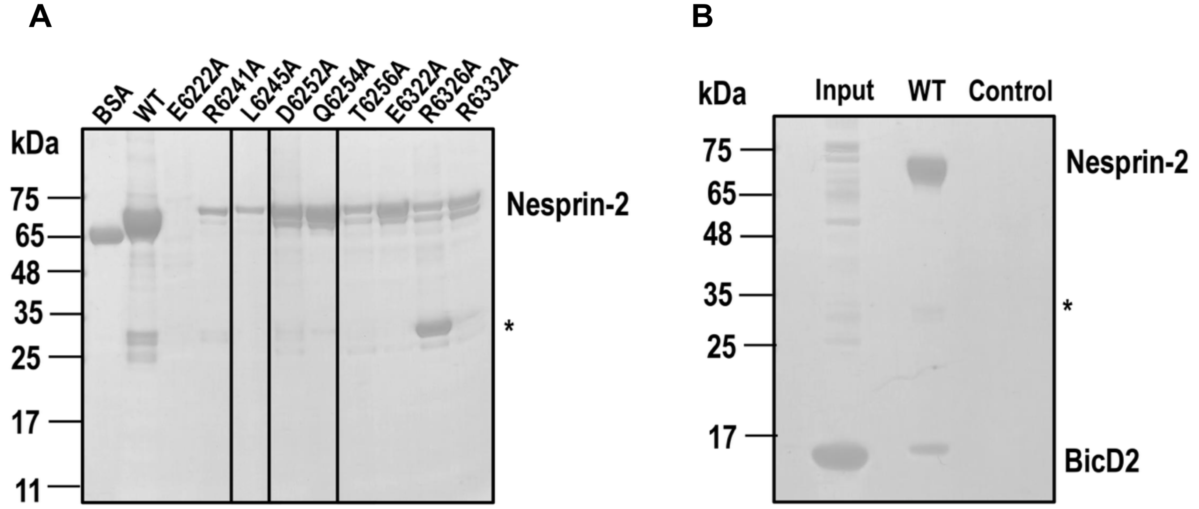


**Figure S9. (A) SDS-PAGE analysis of purified Nesprin-2 mutants that were excluded from the pull-down analysis due to poor solubility.** Proteins were purified from 0.5 l of cells by GST-affinity chromatography and 19 µl of the eluate was analyzed. The positions of the molecular weight standard bands are indicated. Note that the mutants have decreased yields compared to the WT, indicating low solubility. The experiment was performed in three replicates. 3 ug BSA is shown as concentration standard. The asterisk denotes GST. **(B) SDS-PAGE analysis of the negative control for the pull-down assay** (see Figures 2 and 3). The input lane shows purified BicD2 used for the pull-down. The WT Lane shows the elution fraction of a pull-down assay with the WT GST-Nesprin-2 fragment and BicD2-CTD. The control lane shows the negative control, for which BicD2 was added to the empty GST column**.** For a negative control pull-down assay with GST protein and BicD2-CTD see [21]. See also Figure 3.

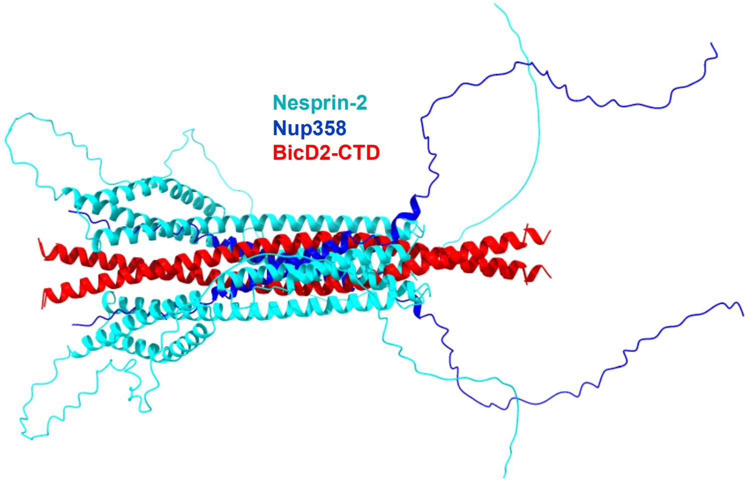


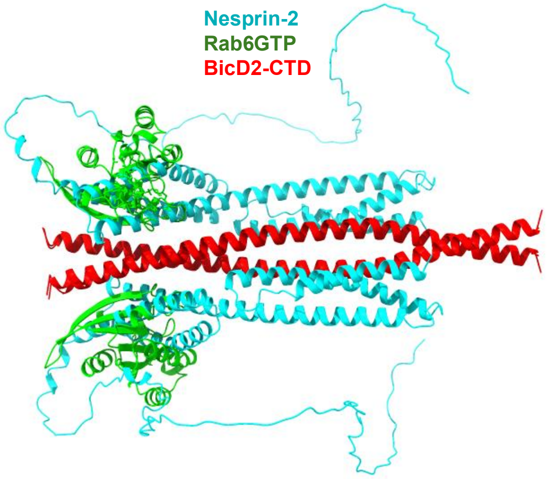


**Comparison of interfaces of BicD2/cargo complexes**

| | Nesprin-2/BicD2 | Nup358/BicD2 | Rab6/BicD2 |
|---|---|---|---|
| **# of contact residues** | 160 | 209 | 96 |
| **Buried surface area Å$^2$** | 2518 | 3715 | 1535 |
| **Secondary structure of the interacting domain (excluding BicD2)** | α-helix | α -helix<br>random coil | α-helix<br>β-sheet<br>random coil |

**Figure S10. The cargo adapters Nesprin-2, Nup358 and Rab6$^{GTP}$ bind to distinct but overlapping sites on the BicD2-CTD.** Least square superimposition of the structural models of Nesprin-2/BicD2 with the structural models of Nup358/BicD2 (top) and Rab6$^{GTP}$/BicD2 (bottom) [22–24]. See also Figure 5.

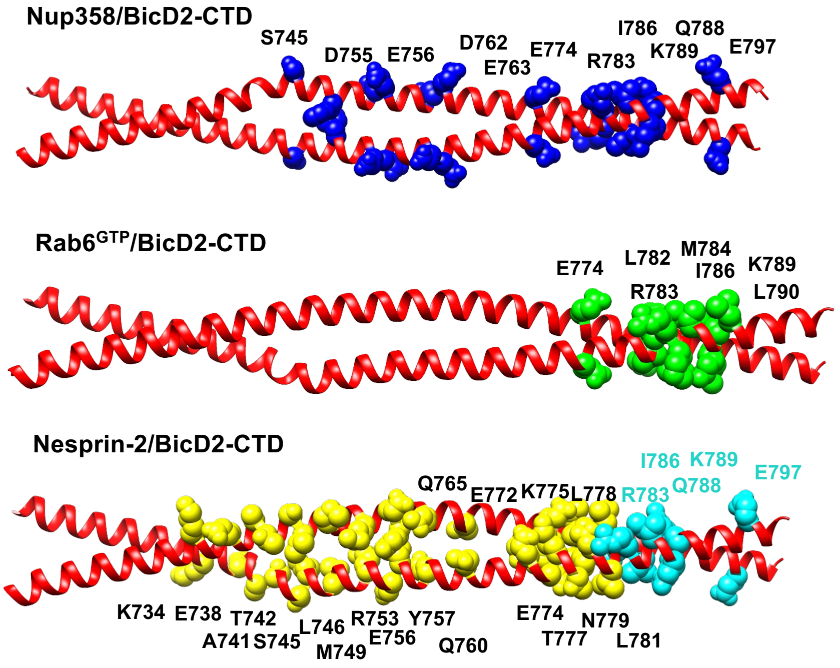


**Figure S11. Comparison of binding sites for Nup358, Nesprin-2 and Rab6$^{GTP}$ on the BicD2-CTD.** The structure of the human BicD2-CTD is shown in red cartoon representation (PDB ID 6OFP)[25]. The contact residues for three cargoes that were verified by mutagenesis and binding assays are highlighted in sphere representation, labeled and colored blue for Nup358 (top panel), green for Rab6$^{GTP}$ (middle panel) and cyan for Nesprin-2 (bottom panel) [21–24,26]. For Nesprin-2, additional BicD2 contact residues from the AlphaFold structural model are highlighted in yellow. See also Figure 5

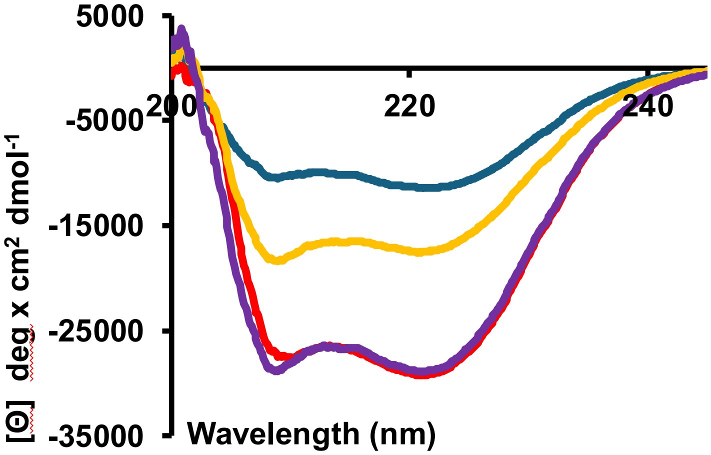


| Molar ellipticity | [Θ] 208 nm (3.5 % error) | [Θ] 222 nm (5% error) |
|---|---|---|
| Nesprin-2/BicD2 complex | -29092 ±1018 | -26224 ±1311 |
| Nesprin-2 + BicD2 | -26224 ±918 | -29092 ±1455 |
| Nesprin-2 | -18147 ±635 | -17402 ±870 |
| BicD2 | -11375 ±398 | -10297 ±515 |

**Secondary structure estimation from the CD spectra with BeStSel [1]**

| Protein | α-helix (%) | Anti-parallel β-sheet (%) | Parallel β-sheet (%) | Turn (%) | Other (%) |
|---|---|---|---|---|---|
| Nesprin-2 | 57.0 | 27.3 | 0 | 15.7 | 0 |
| BicD2 | 21.9 | 24.5 | 1.6 | 16.0 | 36.0 |
| Nesprin-2/BicD2 complex | 49.6 | 28.5 | 15.5 | 6.3 | 0 |
| Nesprin-2 + BicD2 | 49.6 | 28.5 | 15.5 | 6.3 | 0 |

**Figure S12. The minimal Nesprin-2/BicD2-CTD complex has a similar alpha-helical content compared to the individual BicD2-CTD and Nesprin-2 fragments.** A minimal Nesprin-2/BicD2-CTD complex was assembled by mixing the purified Nesprin-2 fragment lacking the intrinsically disordered AD domain (aa 6123-6352, 0.4 mg/ml) with BicD2-CTD (aa 715-804, 0.1 mg/ml). The CD wavelength scan of the assembled complex is shown in purple, overlaid with the CD wavelength scans of the individual proteins (Nesprin-2: yellow; BicD2-CTD: blue) and the sum of the CD wavelength scans of the two individual proteins Nesprin-2 and BicD2-CTD (red). For the CD experiments shown here, the Nesprin-2 fragment lacking the intrinsically disordered AD domain (aa 6123-6352 for sequence see Figure S13) was used, while for the CD experiments in Figure 4 the Nesprin-2 fragment aa 6123-6421 was used. The molar ellipticity [θ] is plotted versus the wavelength. The experiment was repeated three times, a representative experiment is shown. A table lists [θ] at 208 and 222 nm (characteristic wavelength for alpha-helical structures) including experimental error [27]. Note that [θ] at 222 nm of the complex is similar to the sum of the CD wavelength scans of the individual proteins, suggesting a similar alpha-helical content. A secondary structure estimation from the CD spectra was carried out using BeStSel [1].

**Nesprin-2 aa 6123-6352 (single molecule assays, Figure S12)**
MSPILGYWKIKGLVQPTRLLLEYLEEKYEEHLYERDEGDKWRNKKFELGLEFPNLPYYIDGDVK
LTQSMAIIRYIADKHNMLGGCPKERAEISMLEGAVLDIRYGVSRIAYSKDFETLKVDFLSKLPEML
KMFEDRLCHKTYLNGDHVTHPDFMLYDALDVVLYMDPMCLDAFPKLVCFKKRIEAIPQIDKYLK
SSKYIAWPLQGWQATFGGGDHPPKSDLEVLFQGPLGSRLWQKFLDDYSRFEDWLKSAERTAA
CPNSSEVLYTNAKEELKRFEAFQRQIHERLTQLELINKQYRRLARENRTDTASKLKQMVHEGNQ
RWDNLQKRVTAILRRLRYFTNQREEFEGTRESILVWLTEMDLQLTNVEHFSESDAEDKMRQLN
GFQQEITLNTNKIDQLIVFGEQLIQKSEPLDAVLIEDELEELHRYCQEVFGRVSRFHRRLTSHTPG
LDDEKEASENETMDYKDDDDKAGEAPAAAEISGHIVRSPMVGTFYRTPSPDAKAFIEVGQKVN
VGDTLCIVEAMKMMNQIEADKSGTVKAILVESGQPVEFDEPLVVIEGS

**Nesprin-2 aa 6123-6421 (single molecule assays)**
MSPILGYWKIKGLVQPTRLLLEYLEEKYEEHLYERDEGDKWRNKKFELGLEFPNLPYYIDGDVK
LTQSMAIIRYIADKHNMLGGCPKERAEISMLEGAVLDIRYGVSRIAYSKDFETLKVDFLSKLPEML
KMFEDRLCHKTYLNGDHVTHPDFMLYDALDVVLYMDPMCLDAFPKLVCFKKRIEAIPQIDKYLK
SSKYIAWPLQGWQATFGGGDHPPKSDLEVLFQGPLGSRLWQKFLDDYSRFEDWLKSAERTAA
CPNSSEVLYTNAKEELKRFEAFQRQIHERLTQLELINKQYRRLARENRTDTASKLKQMVHEGNQ
RWDNLQKRVTAILRRLRYFTNQREEFEGTRESILVWLTEMDLQLTNVEHFSESDAEDKMRQLN
GFQQEITLNTNKIDQLIVFGEQLIQKSEPLDAVLIEDELEELHRYCQEVFGRVSRFHRRLTSHTPG
LDDEKEASENETDIEDPREIQADSWRKRRESEEPTSPQSLCHLVPPALGHERSGCETPVSVDSI
PLEWDHTGDVGGSSSHETTRRAHSTAHCQMMKQLEDKVEELLSKNYHLENEVARLKKLVGER
MSGDKDCEMKRTTLDSPLGKLELSGCEQGLHRIIFLGKGTSAADAVEVPAPAAVLGGPEPLMQ
ATAWLNAYFHQPEAIEEFPVPALHHPVFQQESFTRQVLWKLLKVVKFGEVISYSHLAALAGNPAA
TAAVKTALSGNPVPILIPCHRVVQGDLDVGGYEGGLAVKEWLLAHEGHRLGKPGLG

**Nesprin-2 aa 6123-6352 (SEC-MALS)**
SPILGYWKIKGLVQPTRLLLEYLEEKYEEHLYERDEGDKWRNKKFELGLEFPNLPYYIDGDVKLT
QSMAIIRYIADKHNMLGGCPKERAEISMLEGAVLDIRYGVSRIAYSKDFETLKVDFLSKLPEMLK
MFEDRLCHKTYLNGDHVTHPDFMLYDALDVVLYMDPMCLDAFPKLVCFKKRIEAIPQIDKYLKS
SKYIAWPLQGWQATFGGGDHPPKSDLEVLFQGPLGSRLWQKFLDDYSRFEDWLKSAERTAAC
PNSSEVLYTNAKEELKRFEAFQRQIHERLTQLELINKQYRRLARENRTDTASKLKQMVHEGNQR
WDNLQKRVTAILRRLRYFTNQREEFEGTRESILVWLTEMDLQLTNVEHFSESDAEDKMRQLNG
FQQEITLNTNKIDQLIVFGEQLIQKSEPLDAVLIEDELEELHRYCQEVFGRVSRFHRRLTSHTPGL
DDEKEASENET

**Figure S13. Protein sequences of the Nesprin-2 fragments used for single molecule assays, SEC-MALS and Figure S12.**

### SUPPORTING MOVIE LEGENDS

**Movie S1. Single-molecule processivity assays of DDB complexes with full length auto-inhibited BicD2 on MTs. The movie is playing at 10× speed.**

**Movie S2. Single-molecule processivity assays of DDBNs complexes with dynein, dynactin, full-length BicD2 and Nesprin-2 fragments on MTs. The movie is playing at 10× speed.**

### SUPPORTING METHODS

#### SEC-MALS

The Nesprin-2 fragment was purified using gel filtration as described under single-molecule experiments, and the BicD2-CTD fragment was purified as described in [28]. Prior to analysis, purified proteins were filtered (0.02 um pore size) and centrifuged in a microcentrifuge (4 °C, 21700g, 30 min). 100 ul sample was injected at room temperature onto a Superdex 200 Increase 10/300 GL column (Cytiva) that was equilibrated with 20 mM HEPES (pH 7.5), 500 mM NaCl, and 0.5 mM TCEP. The column was connected to a Wyatt Technology DAWN 8+ multiangle light scattering detector and an Optilab TrEX refractive index detector. ASTRA 6.1 (Wyatt Technology) was used to determine weight-averaged molar masses.
